# Intrinsic geometry of trial-to-trial variability in primary visual cortex is optimal for robust representation of visual similarity

**DOI:** 10.1101/2025.06.30.662469

**Authors:** Jehyun Kim, Hyeyoung Shin

## Abstract

How neuronal populations construct robust representations of the sensory world despite neural variability remains unclear. Here, we show that trial-to-trial variability in mouse primary visual cortex follows a simple rule: for each stimulus, the mean and variance of spike counts across neurons show a highly stereotyped relationship, with a slope of 1 on a log-log scale. To test how this geometry of trial-to-trial variability affects sensory representations, we numerically manipulated the slope of the log-mean vs. log-variance relationship. We found that the intrinsic geometry of trial-to-trial variability, with slope 1, enables representations of distinct sensory inputs to have minimal overlap while being continuous. At this slope, representational similarity was maximally consistent across neuronal subsets, both within and across mice. By contrast, at slope 0, when variance is uniform across neurons, visual representations were more efficient but less robust. Simulations of recurrent networks, where excitatory and inhibitory neurons formed locally balanced clusters with stronger connections, showed that larger clusters produced steeper slopes and more robust representations. Together, these results suggest that the geometry of trial-to-trial variability mediates a tradeoff between efficiency and robustness, and that the intrinsic geometry is near optimal for robust visual representations.

## Introduction

The neural code involves a tradeoff between efficiency and robustness^1^. In an efficient code, each neuron encodes unique information, which leads to high precision. Conversely, in a robust code, neurons contain redundant information, leading to high reliability. Efficiency vs. robustness is tied to dimensionality: In the extreme case of efficiency, all neurons encode completely non-overlapping information, and the dimensionality is as high as the number of neurons. Pioneering work by Stringer et al.^1^ showed that there is an upper limit of dimensionality allowed for neural representations to be smooth, or differentiable. In other words, if the dimensionality of neural representations is too high, there are instances where infinitesimal differences in sensory inputs lead to large differences in neural representations. Constraining the dimensionality for smoothness results in robust representations. The same study showed that visual representations in mouse primary visual cortex (V1) are at the upper limit of dimensionality^1^, indicating that cortical representations are maximally efficient within the constraint of robustness. However, this work did not look at the role of neural variability.

Variable responses to the same sensory input are a universal characteristic of neocortical neurons, across brain areas and species^2-8^. Whether such trial-to-trial variability is a feature of the neural code, or a bug (so-called ‘noise’), has been intensely debated^9-16^. Variable responses to a given sensory stimulus can cause representations of different stimuli to overlap^17-20^. These overlaps are detrimental to discriminability, precision, and efficiency^20-22^. On the other hand, several studies have suggested that trial-to-trial responses may represent a probability distribution for perceptual inference^23-26^. In this view, overlap between the representations of distinct sensory inputs may allow probabilistic interpretation of an ambiguous input (“What I saw may be A or B”), enabling probabilistic inference based on perceptual similarity. Crucially, given that smoothness is tied to robustness^1^, representational overlap may contribute to robustness by ensuring that representations are continuous across distinct stimuli. Thus, representational overlap may mediate a tradeoff between efficiency and robustness.

Here, we asked how trial-to-trial variability influences the efficiency and robustness of the neural code. We found that spike counts over repeated presentations of the same stimulus had a fixed mean vs. variance relationship across neurons for a given stimulus. Specifically, the log-mean vs. log-variance relationship of spike counts across neurons per stimulus (LMLV_stim_) was highly consistent with slope near 1 across species, visual cortical areas, stimulus categories, familiarity, adaptation and behavioral states. Geometrically, this means that trial-to-trial variability is elliptical and points towards the origin in the neural state space, i.e., the point where all neurons have zero spikes. The neural state space refers to the N-dimensional space of N neurons where each axis indicates the spike count of a neuron during a fixed time window.

To understand the functional role of the highly stereotyped geometry of trial-to-trial variability for the neural code, we manipulated the geometry by parametrically changing the LMLV_stim_ slope among simultaneously recorded mouse V1 neurons. The stimulus manifold, defined by the point cloud of trial-by-trial responses in neural state space, was close to spherical when LMLV_stim_ slope was 0 because all neurons had equal variance. With increasing slope, the stimulus manifolds became increasingly elliptical and lower dimensional. The slope increase also increased the overlap between stimulus manifolds and decreased discriminability. Interestingly, the intrinsic geometry of trial-to-trial variability, with LMLV_stim_ slope near 1, was optimal for consistent representations of visual similarity between distinct subpopulations in mouse V1, both within a mouse and across mice. When the slope decreased below 1, the visual representation space became discontinuous, which was harmful to robustness but beneficial to efficiency. When the slope increased above 1, the geometry of trial-to-trial variability infringed upon the coding dimensions such that both robustness and efficiency were harmed. To understand the neural mechanisms that determine the geometry of trial-to-trial variability, we conducted network simulations. In these simulations, excitatory and inhibitory neurons were divided into locally balanced clusters with stronger connectivity^27^. Cluster granularity shaped the geometry of trial-to-trial variability, such that larger clusters led to more robust representations and steeper LMLV_stim_ slopes. In sum, we found that the geometry of trial-to-trial variability mediates a tradeoff between efficiency and robustness. The highly consistent intrinsic geometry of trial-to-trial variability was suboptimal for efficiency but near optimal for robust visual representations.

## Results

### Stimulus-specific mean vs. variance relationship across neurons is highly stereotyped in visual cortex

We sought to investigate how the neural variability associated with each visual stimulus influences the overall visual representation space. To this end, we examined the geometry of trial-to-trial variability by looking at the relationship between the mean and variance of spike counts during the visual presentation period, across repeated presentations of the same stimulus. We analyzed three extracellular electrophysiology datasets: mouse recordings of visual cortical areas during presentation of natural scene images, static and drifting gratings, and natural movies (Allen Brain Observatory Visual Coding Neuropixels)^28,29^; monkey recordings of primary visual cortex (V1) during presentation of natural scene images^30^; and mouse recordings of V1 during a change detection task with natural scene images (Allen Brain Observatory Visual Behavior Neuropixels)^31^. When we fit a linear regression to the log-mean vs. log-variance relationship across simultaneously recorded neurons in each brain area for each stimulus (LMLV_stim_), we found a highly consistent relationship that was characterized by a slope near 1, and y-intercept near 0.1 (Fig. 1). Note, the slope of a linear regression in the log-log scale corresponds to the exponent in the arithmetic scale; as such, LMLV_stim_ slope of 1 indicates that the mean and variance of spike counts in response to a visual stimulus are linearly related across neurons.

**Fig. 1:**
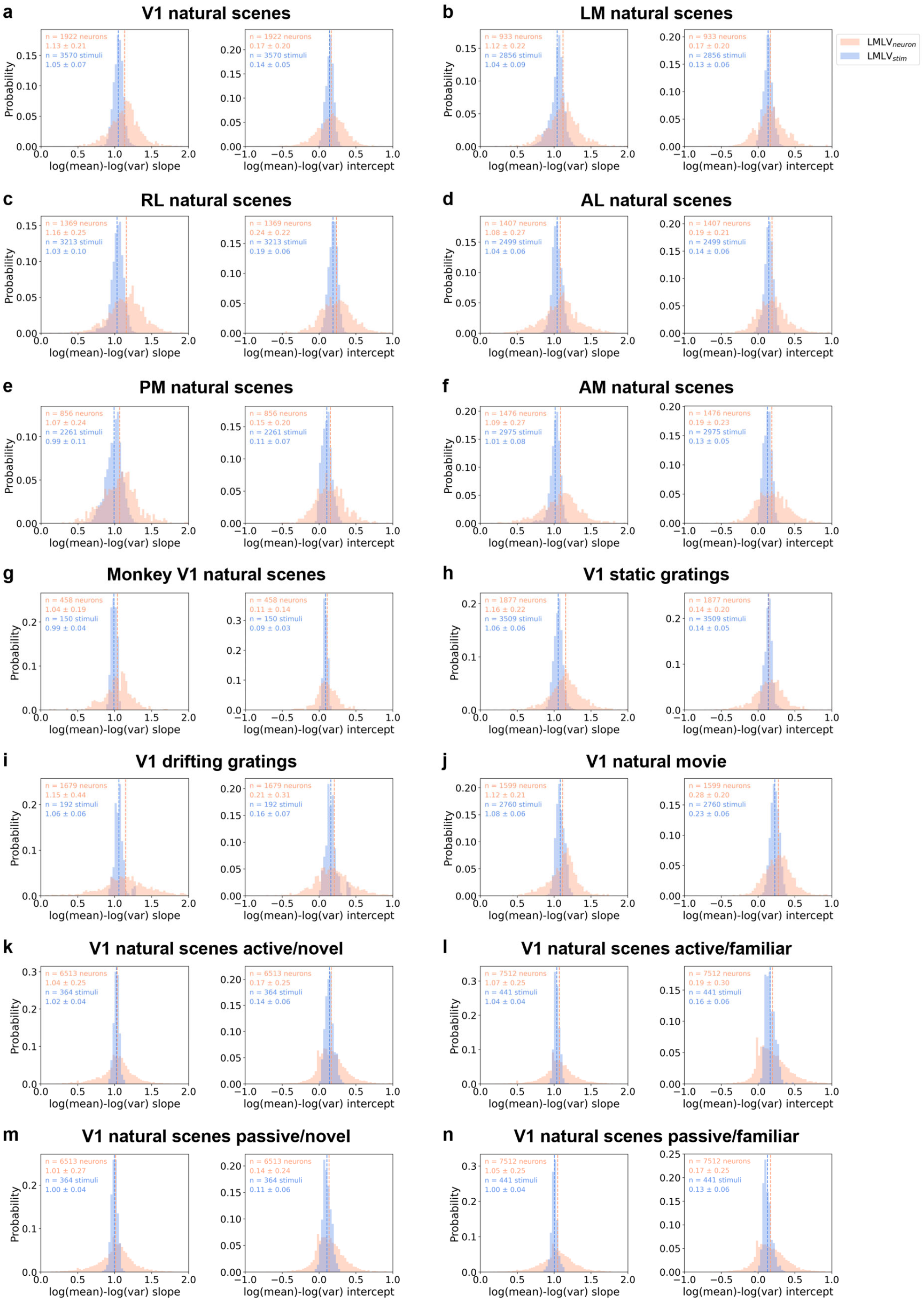
Stimulus-specific mean vs. variance relationships across neurons are universally linear in visual cortex. **a-f** Histograms of log-mean vs. log-variance slopes (*left*) and y-intercepts (*right*) across neurons per stimulus (LMLV_stim_; *blue*) and across stimuli per neuron (LMLV_neuron_; *pink*) for mouse visual cortical neurons’ responses to natural scenes. Spike counts during 0–250 ms after stimulus onset were analyzed, in V1 (**a**), lateromedial (LM, **b**), rostrolateral (RL, **c**), anterolateral (AL, **d**), posteromedial (PM, **e**), and anteromedial (AM, **f**) areas of mouse visual cortex (119 images comprised of 118 natural scenes and a gray image, each repeated ∼50 times in 30 sessions of 30 mice). **g** Same histograms for monkey V1 neurons’ responses to natural scenes. Spike counts during 40–160 ms after stimulus onset were analyzed (75 natural scenes were repeated ∼40 times in 32 sessions of 2 monkeys). Neurons were pooled in each monkey for LMLV_stim_ (238 single units in 15 sessions of monkey M1, 220 single units in 17 sessions of monkey M2). **h** Same histograms for mouse V1 neurons’ responses to static gratings. Spike counts during 0–250 ms after stimulus onset were analyzed (121 images comprised of 120 static gratings and a gray image, each repeated ∼50 times in 29 sessions of 29 mice). **i** Same histograms for mouse V1 neurons’ responses to drifting gratings. Spike counts during 0–250 ms after stimulus onset were analyzed (8 drifting gratings repeated 75 times in 24 sessions of 24 mice). **j** Same histograms for mouse V1 neurons’ responses to a natural movie. Spike counts were calculated for 250 ms bins of a 30 s movie clip, resulting in 120 consecutive windows (‘stimuli’). The movie was repeated 60 times in 23 sessions of 23 mice. **k-n** Same histograms for mouse V1 neurons’ responses to natural scenes during a change detection task (‘active’), and subsequent passive replay (‘passive’), for familiar and novel stimuli. Spike counts during 0–250 ms after stimulus onset were used. In familiar sessions, 8 familiar images used during training were shown (49 sessions of 49 mice). In novel sessions, 6 novel images and 2 familiar images were shown, and only the novel image trials were analyzed (52 sessions of 52 mice). Each image was repeated ∼580 times per session, with ∼3% probability of omission (gray image; analyzed as a separate ‘stimulus’). **a-n** Mean ± standard deviation was indicated in each plot, across either stimuli pooled across sessions (LMLV_stim_) or neurons pooled across sessions (LMLV_neuron_). The dotted lines indicate the mean of each condition.

Remarkably, LMLV_stim_ was highly consistent among natural scene images not only across visual cortical areas of mice (Fig. 1a-f), but also in monkeys (Fig. 1g). Further, LMLV_stim_ was consistent regardless of stimulus category (Fig. 1h-j), novelty vs. familiarity, and active vs. passive engagement in a visual task (Fig. 1k-n). Notably, the order of stimulus presentation was randomized in all datasets except for the Allen Brain Observatory Visual Behavior Neuropixels dataset, where stimulus presentations were repeated four or more times at regular intervals of 750 ms (250 ms image presentation and 500 ms gray screen)^31,32^. Despite the presence of adaptation in this data^33^, the LMLV_stim_ was highly stereotyped with slope concentrated around 1 (Fig. 1k-n).

To further investigate these mean vs. variance relationships, we focused on mouse V1 responses to natural scenes, from the Allen Brain Observatory Visual Coding Neuropixels dataset (Fig. 1a)^28,29^. In this dataset, awake head-fixed mice viewed repeated presentations of natural scene images (119 images, consisting of 118 natural scenes and 1 gray screen image, presented for 250 ms with no intervening period, repeated 50 times each in a randomized order; 14–110 single units per session, n = 30 sessions from 30 mice).

The log-mean vs. log-variance relationship across neurons per stimulus (LMLV_stim_) was more linear and more narrowly distributed compared to the supralinear relationship across stimuli per neuron (LMLV_neuron_; Fig. 1). LMLV_stim_ and LMLV_neuron_ describe mean vs. variance relationships at the population level and single-neuron level, respectively. The supralinear mean vs. variance relationship for single neurons has been studied extensively in the literature ^4,24,34-36^. Prior work by Goris et al.^4^ suggested that mean vs. variance relationship per neuron can be explained by the Modulated Poisson model, where a Poisson spike process is modulated by a multiplicative gain that fluctuates independently of a stimulus. When the multiplicative gain fluctuates with a gamma distribution, the spike count distribution follows a negative binomial distribution NB(*r, p*) where the parameter *r* is the inverse of the gain variance 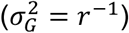. Because multiplicative gain fluctuations are independent of the stimulus, the Modulated Poisson model predicts that spike count distributions across stimuli per neuron follow negative binomial distributions with a fixed *r* parameter. Indeed, when we fit negative binomials to spike count distributions for each neuron and stimulus in the mouse V1 natural scenes data, we found that gain variance was largely consistent across stimuli per neuron, but heterogeneous across neurons per stimulus (Supplementary Fig. 1a, b). In addition, 10-fold cross-validated negative log likelihood, Akaike information criterion (AIC), and root mean squared error (RMSE) were not consistently higher nor lower for the Modulated Poisson model compared to linear fits of LMLV_neuron_, indicating that both models are equally valid for describing the mean vs. variance relationship of each neuron (Supplementary Fig. 1f). In summary, LMLV_neuron_ was consistent with the Modulated Poisson model^4^, implicating stimulus-independent multiplicative gain as a source of trial-to-trial variability. In contrast, the population-level homogeneity of LMLV_stim_ has not been systematically investigated. As such, we focused on the population geometry of trial-to-trial variability, LMLV_stim_, in the remainder of this paper.

The LMLV_stim_ was highly stereotyped across stimuli in various categories, suggesting that LMLV_stim_ is intrinsically shaped by the cortical network. If so, the mean vs. variance relationship across neurons in the absence of a stimulus should also be highly stereotyped. To test this prediction, we analyzed the log-mean vs. log-variance of spike counts across V1 neurons for prolonged spontaneous periods (LMLV_spon_; 30-minute gray-screen period was divided into non-overlapping bins of 250 ms, and 50 bins were subsampled). We found that LMLV_spon_ also has a slope near 1 (mean ± standard deviation 1.07 ± 0.08, Supplementary Fig. 2a). However, the y-intercept was higher for LMLV_spon_ than LMLV_stim_, consistent with stimulus driven variability quenching^37^ (mean ± standard deviation 0.25 ± 0.06 for LMLV_spon_, 0.14 ± 0.05 for LMLV_stim_ natural scenes excluding the interleaved gray-screen trials, Supplementary Fig. 2b). Interestingly, gray-screen trials interleaved among natural scene trials had slopes and y-intercepts that were indistinct from the rest of the natural scene images. As such, interleaved gray-screen trials and prolonged gray-screen periods (LMLV_spon_) had similar slopes but significantly different y-intercepts (Supplementary Fig. 2a, b). This may be because visual offset transiently drives feedforward inputs to visual cortex.

LMLV_stim_ slopes and y-intercepts were slightly but significantly modulated by locomotion and pupil size, having lower values when locomoting (Supplementary Fig. 2c, d) and when pupils were dilated (Supplementary Fig. 2e, f). Although this modulation was significant, the effect was small, such that the slope remained near 1 across a range of behavioral states.

In addition, we checked whether our results could be reproduced in two-photon calcium imaging data (Supplementary Fig. 3). We utilized the Allen Brain Observatory Visual Coding two-photon dataset^38,39^, where mouse V1 calcium activity was recorded during presentation of the exact same natural scene images. We found that neither dF/F nor deconvolved ‘spikes’^40^ of GCaMP6f activity recapitulated the mean vs. variance relationship of spike counts per neuron (LMLV_neuron_), nor per stimulus (LMLV_stim_). These results indicate that spike counts cannot be accurately measured with two-photon calcium imaging.

### Geometry of trial-to-trial variability governs representational overlap

We have seen that the geometry of trial-to-trial variability is highly stereotyped across a plethora of conditions, characterized by LMLV_stim_ slope of 1. To investigate the functional implication of this phenomenon, we created pseudo-data where the linear regression of LMLV_stim_ follows a new slope. The y-intercept was set to keep the total log-variance constant (Fig. 2b). Based on the target LMLV_stim_ slope and y-intercept, we calculated the target variance for each neuron and stimulus and rescaled the residuals in the trial-by-trial responses by multiplying a constant (Fig. 2d; see Methods). This manipulation preserves the mean spike count for each neuron and stimulus, total log-variance across neurons per stimulus, and pairwise noise correlations for each neuronal pair and stimulus.

**Fig. 2:**
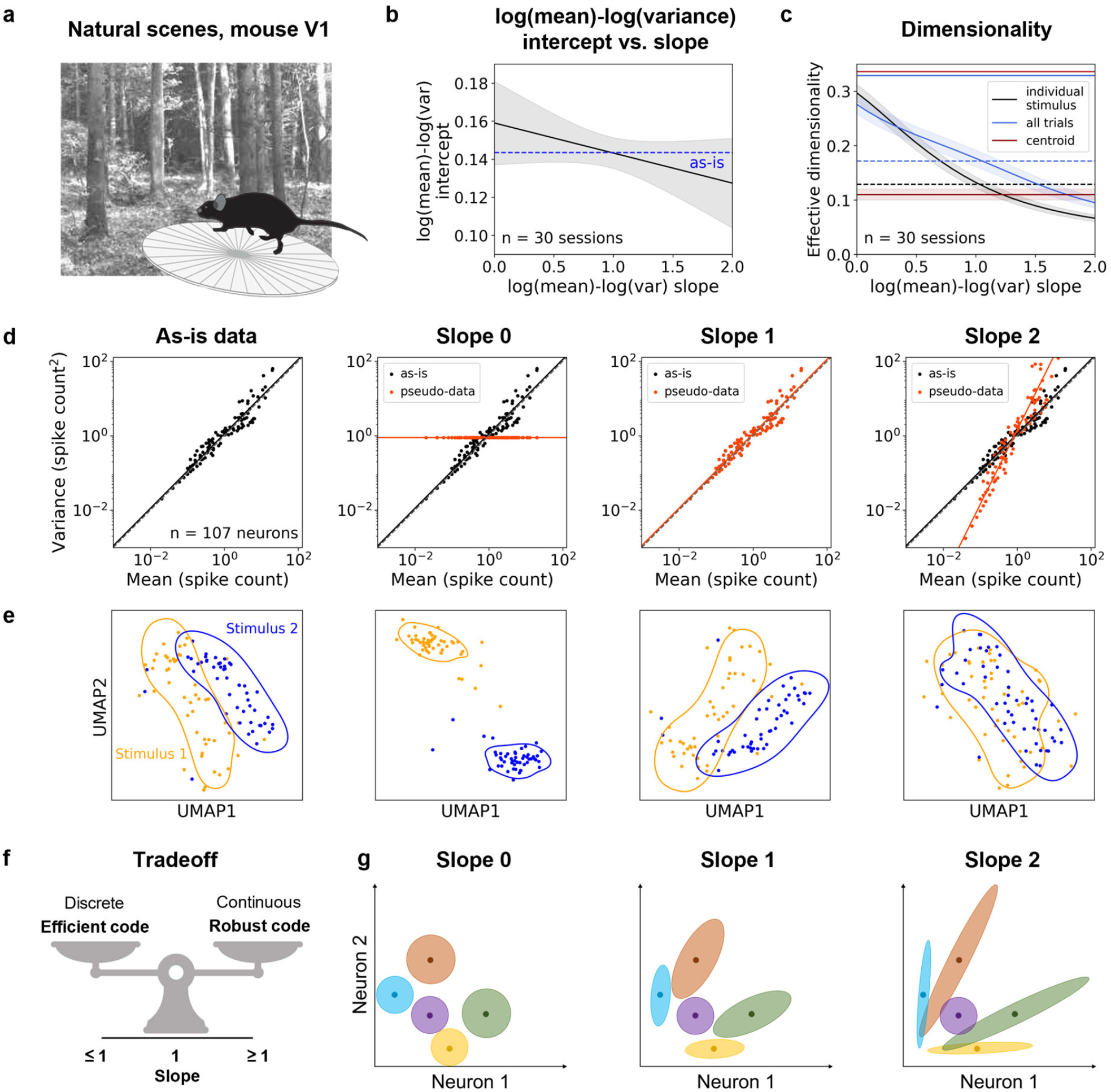
Geometry of trial-to-trial variability governs representational overlap. **a** Schematic of the main dataset: 119 images (118 natural scenes and a gray image) were presented for 250 ms with no intervening gray screen period, each repeated 50 times. **b** LMLV_stim_ y-intercept after manipulation of LMLV_stim_ slope. For each target LMLV_stim_ slope from 0 to 2, target y-intercept was set to keep the total log-variance constant. **c** Effective dimensionality of individual stimulus manifolds (*black*), all stimulus manifolds (all trials; *blue*), and the centroids of all stimulus manifolds (*red*). Effective dimensionality of individual stimulus manifolds was averaged across stimuli within each session. The dotted lines indicate the mean of as-is data for each condition. **b, c** Mean ± standard error mean (SEM) across sessions was plotted as a function of LMLV_stim_ slopes. The colored lines on top indicate slopes that are significantly different from the as-is data (p < 0.05, two-sided Wilcoxon signed-rank test across sessions, Holm-Bonferroni corrected). In **b**, there was no slope significantly different from as-is data. **d** LMLV_stim_ scatterplots and slope manipulation for an example natural scene image in an example session. Each *black* dot corresponds to a single neuron in the as-is data, and the *black* line is the linear regression fit of the *black* dots. The *orange* dots and lines show slope-manipulated data, where only the variance was changed. **e** UMAP embeddings of 2 example natural scene manifolds in an example session. Each color denotes one stimulus manifold, and each dot corresponds to a trial (50 trials per stimulus). The contour of each manifold was obtained through kernel density estimation. **f** Schematic illustration of a tradeoff between efficiency and robustness of the neural code mediated by the LMLV_stim_ slope. **g** Schematic illustration of stimulus manifolds across LMLV_stim_ slopes. The bold points indicate the centroids of the 5 stimulus manifolds.

The LMLV_stim_ slope increase resulted in lower effective dimensionality (Fig. 2c), leading to corresponding changes in the stimulus manifold geometry. Since mean spike counts do not change with slope manipulation, the centroid of each stimulus manifold in the neural state space stays in place. To examine how the slope manipulation changes the geometry of stimulus manifolds in the neural state space, we applied a nonlinear dimensionality reduction method called UMAP^41^ (Fig. 2e). When the slope is 0, the variance is the same across all neurons. Hence, the stimulus manifold is nearly spherical, deviating only because of noise correlations (Fig. 2e, g). Neighboring stimulus manifolds appear discontinuous with each other, prohibiting smooth transitions between them. As the slope increases, neurons with larger means have larger variances, leading to increasingly elliptical stimulus manifolds that point toward the origin in the neural state space (Fig. 2e, g). The overlap between stimulus manifolds also increases. At slope 2, the stimulus manifolds are hard to discriminate due to excessive overlap. The stimulus manifolds appear to strike a balance when the slope was near 1, such that they are continuous with minimal overlap in the original ‘as-is’ data (Fig. 2f).

To better understand the functional significance of a continuous representation, we examined the neural representation of a natural movie in the same dataset. The movie clip did not contain any abrupt transitions between consecutive frames. Since the visual inputs were continuous in time, the neural trajectory should also be continuous in time. As expected, the neural trajectory during natural movie presentation was continuous in the UMAP (60 repetitions of ‘natural movie one’ in an example session with 85 V1 neurons; Supplementary Fig. 4). The log-mean vs. log-variance of spike counts during a designated movie segment showed a linear relationship across neurons with slope near 1 (Fig. 1j), as in the natural scenes data. When we manipulated the LMLV_stim_ slope, the neural trajectory broke into several disjoint segments at slope 0. At slope 2, the trajectory was continuous, but the time information was blurred. From this example, we can gather the intuition that visual representations involve a tradeoff between maintaining continuity and avoiding excessive overlap. This tradeoff appears to be balanced at slope 1.

### Intrinsic geometry of trial-to-trial variability enables maximal discriminability within the constraint of a continuous neural code

We have observed via dimensionality reduction that neighboring stimulus manifolds that were discontinuous at LMLV_stim_ slope 0 became continuous at slope 1 and showed excessive overlap at slope 2. This implies that the overlap between stimulus manifolds generally increase with LMLV_stim_ slope. To formalize this observation, we sought to quantify overlap and continuity in the neural state space, without relying on dimensionality reduction. First, to quantify overlap, we defined the overlap percentage of stimulus manifold B relative to A as the fraction of points in B that were closer to a point in A than the bottom 5^th^ percentile of pairwise Euclidean distances within A (Fig. 3a *left*)^42^. We averaged the overlap percentage across all stimulus pairs in each session (Fig. 3a *right*). We found that overlap percentage increases with slope, in good agreement with our observations of the dimensionality-reduced visual representation space (Fig. 2e).

**Fig. 3:**
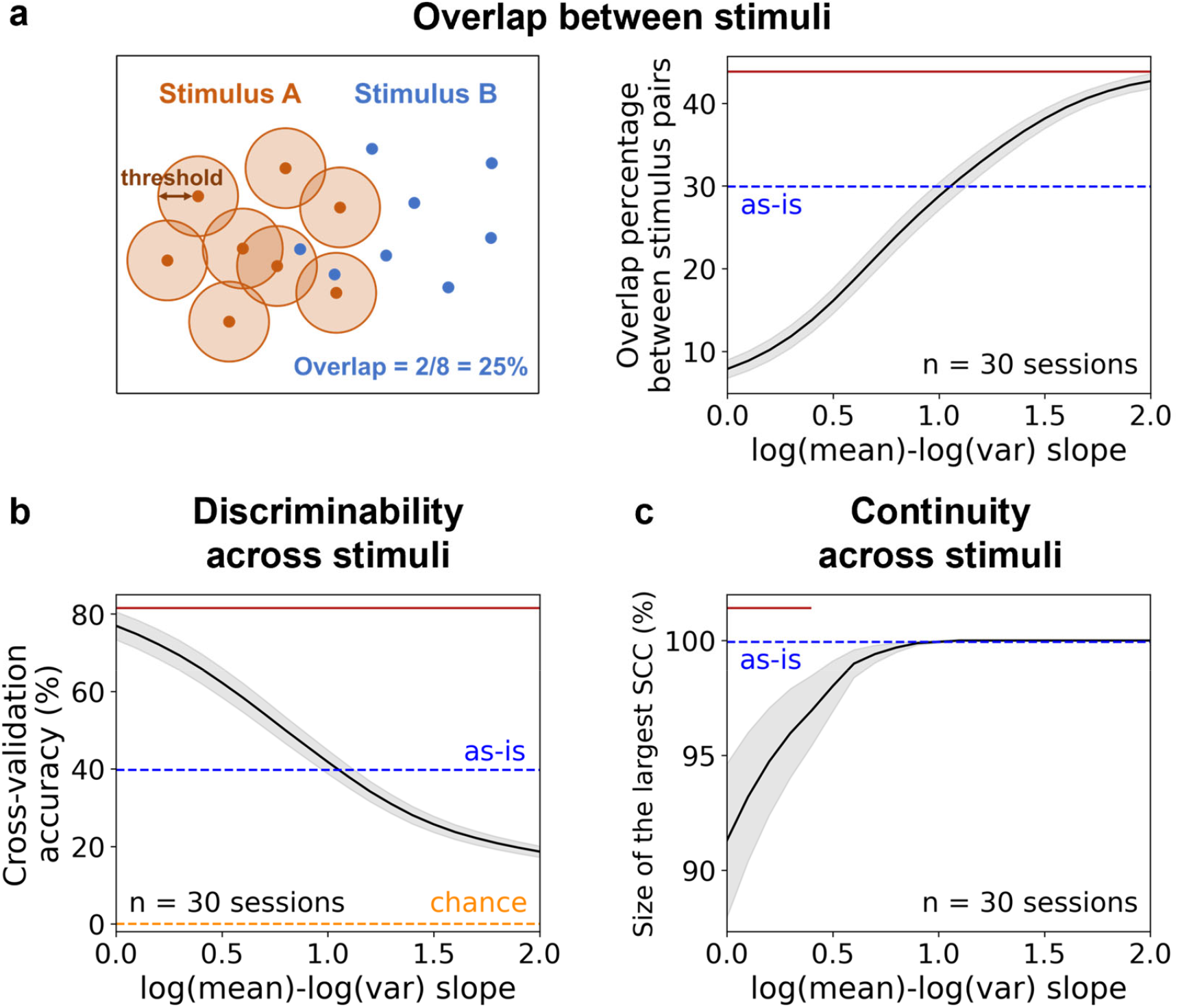
Intrinsic geometry of trial-to-trial variability enables maximal discriminability within the constraint of a continuous neural code. **a** Overlap percentage between stimulus manifolds. *Left*: Schematic illustration of overlap percentage of stimulus manifold B relative to A. *Right*: Overlap percentage as a function of LMLV_stim_ slope. Percentage of overlap was calculated for every pair of stimulus manifolds and averaged within each session. **b** 10-fold cross-validation classification accuracy of linear support vector machine (SVM) decoder. *Orange* dotted line indicates chance level (1/119 ≈ 0.008). **c** Size of the largest strongly connected component (SCC) in the stimulus manifold graph. The percentage of stimulus manifolds in the largest SCC was plotted. **a-c** Mean ± SEM across sessions was plotted as a function of LMLV_stim_ slopes. The *red* lines on top indicate slopes that are significantly different from the as-is data (p < 0.05, two-sided Wilcoxon signed-rank test across sessions, Holm-Bonferroni corrected).

Overlap in neural representations impairs discriminability^43^. Thus, discriminability should decrease with LMLV_stim_ slope increase. To test this prediction, we leveraged linear support vector machines (SVM) to decode image identity from trial-by-trial neural activity vectors, using 10-fold cross-validation (Fig. 3b). As expected, cross-validation accuracy decreased monotonically as a function of the LMLV_stim_ slope. Decoding accuracy is an indicator of neural information, and in turn, coding efficiency^44^. Hence, the intrinsic geometry of trial-to-trial variability normally observed in visual cortex is suboptimal for coding efficiency.

Despite the suboptimal coding efficiency, the visual cortex consistently operates at a LMLV_stim_ slope near 1, suggesting that the intrinsic geometry of trial-to-trial variability may be beneficial in other ways. We observed that the visual representation space appears discontinuous at slope 0 whereas the original, as-is data has a continuous representation (Fig. 2e). This result suggests that the representational space loses continuity at slope 0, thereby compromising smoothness and robustness. Thus, we hypothesized that the intrinsic geometry of trial-to-trial variability enables maximal discrimination within the constraint of a continuous neural code. If so, slope 1 should be the minimum slope that satisfies the continuity constraint.

To determine whether the visual representation space was continuous, we defined a graph of stimulus manifolds. Each node represents a stimulus manifold, and a positive overlap percentage defines a directed connection between the pair of stimulus manifolds. The visual representation space is continuous if all stimulus manifolds are interconnected. Analyzing the size of the largest strongly connected component (SCC) confirmed our prediction, showing that the visual representation space became continuous starting around LMLV_stim_ slope 1 (Fig. 3c).

To maximize discriminability within the constraint of a continuous representation space, we hypothesized that stimulus manifolds should be shaped such that less variance is allocated towards nearby stimuli and more variance towards faraway stimuli. To test this hypothesis, we examined the correlation between the geodesic distance and the orthogonal variance between pairs of stimulus manifolds (Supplementary Fig. 5a-d). The orthogonal variance quantifies the amount of variance in one stimulus manifold directed towards the centroid of a different manifold (Supplementary Fig. 5a)^45^. As predicted, we found that the geodesic distance between the centroids of a pair of stimulus manifolds had a positive correlation with the orthogonal variance (Supplementary Fig. 5b, c). When manipulating the LMLV_stim_ slope, the correlation between geodesic distance and orthogonal variance was highest near the slope of 1 (Supplementary Fig. 5d). These results imply that slope 1 is the most balanced structure of variability in that it optimizes the tradeoff between minimizing overlap and maintaining continuity, which respectively contribute to coding efficiency and robustness.

In sum, we found that while discriminability was highest at LMLV_stim_ slope of 0, the visual representation space was discontinuous. This implies that while visual representations are maximally informative and therefore most efficient at slope 0 (Fig. 3b, Supplementary Fig. 6a), they may be less robust. Thus, we hypothesized that the geometry of trial-to-trial variability mediates the tradeoff between the efficiency and robustness of the neural code, and that at slope 1, visual representations can be both efficient and robust (Fig. 2f).

### Intrinsic geometry of trial-to-trial variability is optimal for consistent representations between distinct neuronal subsets in V1 within a mouse

Next, we sought to elucidate how the LMLV_stim_ slope influences the robustness of neural representations. In a robust code, neurons encode redundant information^46,47^, such that representations by distinct subsets of neurons are consistent. We hypothesized that distinct subsets of a neural population would have less consistent representations when the visual representation space is discontinuous. Thus, when LMLV_stim_ slope increases from 0 to 1, we predicted a tradeoff between a decrease in efficiency and an increase in robustness. Overlap between representations continues to increase with slope, even after becoming globally continuous at slope 1. Thus, we further predicted that LMLV_stim_ slope increase beyond 1 would generate excessive overlap, which would degrade both efficiency and robustness of neural representations.

To test whether distinct subsets of neurons carrying redundant spatial information produced consistent representations, we randomly divided V1 neurons in each session into two halves. We defined overlap consistency as the Spearman correlation between overlap percentages of the two subsets in each session. Overlap consistency peaked near slope 1 (Fig. 4a), suggesting that visual representations are most consistent between neuronal subsets when the LMLV_stim_ slope is near 1.

**Fig. 4:**
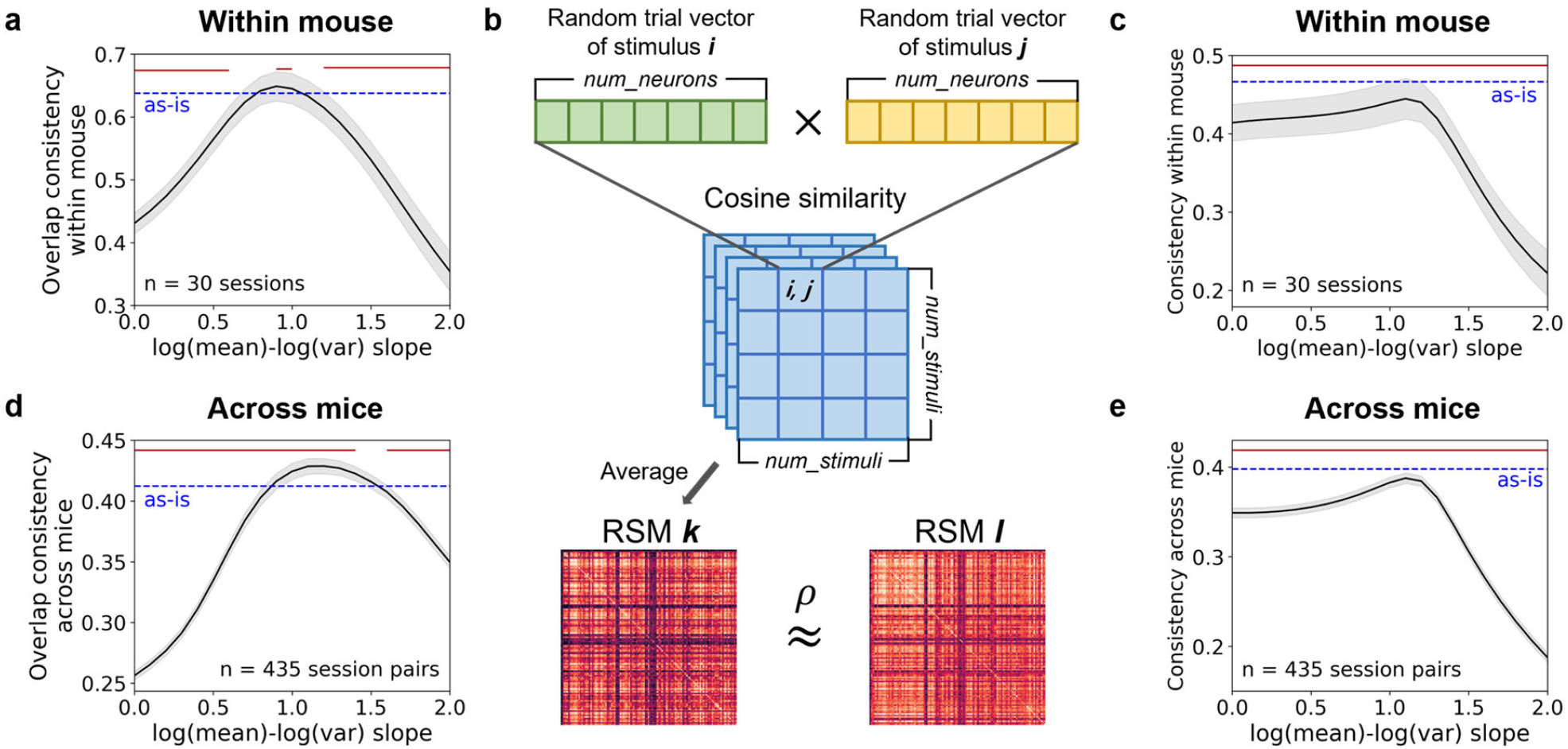
Intrinsic geometry of trial-to-trial variability is optimal for consistent representations in V1 between different neuronal subsets, both within and across mice. **a** Overlap consistency between distinct neuronal subsets within each mouse. **b** Schematic of representational similarity analysis (RSA). Representational similarity matrices (RSM) of two example sessions (mice) are shown at the bottom. **c** Consistency of RSMs between distinct neuronal subsets within each mouse. **d** Overlap consistency between pairs of mice. **e** Consistency of RSMs between pairs of mice. **a, c, d, e** Mean ± SEM across sessions (**a, c**) or across session pairs (**d, e**) was plotted as a function of LMLV_stim_ slopes. The *red* lines on top indicate slopes that are significantly different from the as-is data (p < 0.05, two-sided Wilcoxon signed-rank test across sessions or session pairs, Holm-Bonferroni corrected).

To confirm this effect with a more widely used metric, we leveraged the representational similarity analysis (RSA; Fig. 4b)^48^. Again, we divided V1 neurons in each session into two halves. To incorporate trial-by-trial variability in RSA, we calculated the cosine similarity between trial-by-trial neural activity vectors. We randomly selected a pair of trials for each stimulus pair to construct the #stimuli-by-#stimuli cosine similarity matrix. We repeated this process 1,225 (_50_C_2_) times and averaged the cosine similarity matrices to acquire a representational similarity matrix (RSM) for each neuronal subset. For the pair of RSMs in each session, we calculated the Spearman correlation to assess their consistency. When we applied RSA across parametric variations in the LMLV_stim_ slope, we found that the RSMs in distinct neuronal subsets within V1 were most consistent with each other around slope 1 (Fig. 4c). Taken together, these results suggest that the visual representation subspaces are most consistent when LMLV_stim_ slope is around 1. Thus, the intrinsic geometry of trial-to-trial variability in V1 is optimal for robust representation of similarity.

### Intrinsic geometry of trial-to-trial variability is optimal for consistent representations in V1 between different mice

We have established that the intrinsic geometry of trial-to-trial variability allows distinct neuronal subsets within a mouse V1 to have consistent representations. We asked whether the geometry of trial-to-trial variability might also influence representational consistency across different mice. Consistent visual representation across different mice would imply that different individuals can learn a common model of the sensory world despite different experiences.

We hypothesized that deviation from LMLV_stim_ slope 1 would impair the representational consistency and overlap consistency between different brains, similar to those between distinct neuronal subsets within the same brain. To test this, we applied the same analyses as Fig. 4a, c to pairs of mice (note, there is a one-to-one correspondence between mice and sessions in this dataset). We found that both overlap consistency and the consistency of RSMs peaked around LMLV_stim_ slope of 1 (Fig. 4d, e). Moreover, leave-one-mouse-out analyses also showed highest representational consistency around LMLV_stim_ slope 1 (Supplementary Fig. 6b).

Thus, visual representations were most consistent around slope 1, both within and across mice (Fig. 4). These results held when excluding low-firing rate neurons (Supplementary Fig. 7a-e), subsampling trials (Supplementary Fig. 7f-k), changing the spike count time window (Supplementary Fig. 8), or excluding neurons whose receptive fields were not on the screen (Supplementary Fig. 9a-f).

Additionally, we examined representational consistency between neuronal subsets with distinct receptive fields (Supplementary Fig. 9g-l). We only included sessions where each half of the screen had 10 or more neurons, which substantially reduced the number of sessions that we could analyze (11/30 sessions for left vs. right, 5/30 sessions for top vs. bottom). Therefore, in many cases, the slope-manipulated pseudo-data were not significantly different from the as-is data. Nevertheless, representational consistency across mice between left vs. right sides of the screen, which had the highest number of samples (n = 55 session pairs), showed significant differences (Supplementary Fig. 9i). As in the representational consistency between random subsets (Fig. 4e), the peak consistency was observed near slope 1. Thus, the intrinsic geometry of trial-to-trial variability may be beneficial for consistent representation of visual similarity even between subpopulations with distinct spatial information.

Our slope manipulation method did not change pairwise noise correlations (Supplementary Fig. 10a), but it did change population-wise noise correlations, defined as the fraction of variance explained by the first principal component of each stimulus manifold^49^ (Supplementary Fig. 10b). When we destroyed pairwise noise correlations by shuffling the order of trials within each stimulus for each neuron, the dimensionality increased (Supplementary Fig. 10c, d). The decoding accuracy also increased (Supplementary Fig. 10e), consistent with prior literature^12,50^. However, the effect of pairwise noise correlations on decoding accuracy was small relative to the effect of LMLV_stim_ slope manipulations. LMLV_stim_ slope-dependent effects were largely consistent with and without shuffling: Dimensionality and decoding accuracy decreased with increasing slope (Supplementary Fig. 10c-e), visual representation space became continuous starting around slope 1 (Supplementary Fig. 10f), and representational consistency within a mouse peaked around slope 1 (Supplementary Fig. 10g). In comparison, the representational consistency across mice peaked around slope 1.3 in the trial-shuffled pseudo-data (Supplementary Fig. 10h). This suggests that pairwise noise correlations may slightly shift the balance between efficiency and robustness.

The slope manipulation method based on residual rescaling guarantees that total log-variance and pairwise noise correlations are preserved, but it does not guarantee non-negative integer ‘spike counts’ in the slope-manipulated pseudo-data. Thus, we devised a slope manipulation method based on spike count sampling, where spike counts are sampled from discrete probability distributions (Supplementary Fig. 11). Specifically, we calculated the target mean and variance of the slope-manipulated pseudo-data for each stimulus and neuron combination, as before. Then, depending on the target dispersion of the pseudo-data, we sampled spike counts from negative binomial, binomial, or Poisson distributions for overdispersed, underdispersed, and approximately equidispersed data, respectively (var ≥ 1.05×mean, var ≤ 0.95×mean, and 0.95×mean < var < 1.05×mean; Supplementary Fig. 11a, b). The resulting LMLV_stim_ slopes had a smaller range than the target slope (Supplementary Fig. 11c), and the total log-variance increased with LMLV_stim_ slope (Supplementary Fig. 11d). Nevertheless, we could test whether the LMLV_stim_ slope-dependent effects were consistent with the residual rescaling method. Since pairwise noise correlations were ignored during spike count sampling, we compared the sampled pseudo-data with the residual-rescaled pseudo-data in which pairwise noise correlations were destroyed via trial shuffling (Supplementary Fig. 10b, d-h). These two methods of slope manipulation showed largely consistent slope-dependent trends: Dimensionality and decoding accuracy decreased with slope, and representational consistency within and across mice peaked at a slope greater than 1 (Supplementary Fig. 11e-h). These results confirm that our LMLV_stim_ slope manipulation effects can be reproduced with non-negative integer spike counts.

### Geometry of trial-to-trial variability mediates efficiency-robustness tradeoff in higher visual areas and in monkeys

We have seen that the intrinsic geometry of trial-to-trial variability is suboptimal for efficient discrimination (Fig. 3b), but optimal for robust representations in V1, both within and across mice (Fig. 4). Next, we asked whether these results generalized to other visual cortical areas (Supplementary Fig. 12). Like V1, dimensionality and decoding accuracy decreased with increasing LMLV_stim_ slope in higher visual areas (Supplementary Fig. 12a, b). However, representational consistency within a mouse was not significantly lower than as-is at LMLV_stim_ slopes less than 1 (Supplementary Fig. 12c). On the other hand, representational consistency across mice peaked near slope 1 in several higher visual areas, including LM, AL and PM, similar to V1 (Supplementary Fig. 12d). In short, LMLV_stim_ slope-dependent trends in higher visual areas were similar to V1 for dimensionality, decoding accuracy and representational consistency across mice.

Further, we examined representational consistency between visual cortical areas, both within and across mice (Supplementary Fig. 13). Within a mouse, representational consistency was higher within an area than between areas (Supplementary Fig. 13a). Across mice, representational consistency was similar within and between areas (Supplementary Fig. 13b). In general, representational consistency between visual cortical areas peaked between LMLV_stim_ slope 1–1.5, both within and across mice (Supplementary Fig. 13c, d).

Next, we asked whether LMLV_stim_ slope mediates the efficiency vs. robustness tradeoff in other stimulus categories, such as drifting gratings (Supplementary Fig. 14; 8 drifting gratings in 4 directions and 2 contrasts, 75 repeats; spike counts from 0–250 ms relative to visual onset). Like natural scenes, drifting gratings also had LMLV_stim_ slopes that were consistently near 1 (Fig. 1i), and the dimensionality and decoding accuracy decreased with LMLV_stim_ slope (Supplementary Fig. 14a-c). However, representational consistency within and between mice were not significantly different from as-is at LMLV_stim_ slopes below 1 (Supplementary Fig. 14d, e). When we examined the RSMs, we observed prominent diagonals at slope 0 (Supplementary Fig. 14f). The diagonal terms quantify trial-to-trial reliability per stimulus, which is generally correlated with cross-validation decoding accuracy. To isolate the between-stimuli effects, we excluded the diagonal terms (Supplementary Fig. 14g, h). Without the diagonals, representational consistency across mice was significantly lower at slopes below 1 than as-is (Supplementary Fig. 14h). These results suggest that the geometry of trial-to-trial variability generally mediate the balance between efficient vs. robust representation of visual similarity, but that the balance point may slightly depend on the stimulus category.

Lastly, we wondered whether the geometry of trial-to-trial variability also mediated efficiency vs. robustness tradeoff in monkeys (Supplementary Fig. 15). To address this, we analyzed publicly available extracellular electrophysiology recordings of monkey V1, during presentation of 75 natural scene images repeated ∼40 times in randomized order^30^. This dataset had fewer simultaneously recorded V1 units than the mouse data (14.3 ± 7.6 V1 units per session in monkeys vs. 63.0 ± 24.8 in mice; mean ± standard deviation across sessions). To gain statistical power, we created 15 pseudo-sessions of 110 neurons in each monkey. As these pseudo-sessions contained neurons that were not simultaneously recorded, we destroyed pairwise noise correlations altogether by randomizing the trial order of each image for each neuron. When comparing with the trial-shuffled pseudo-data in mice, we found similar LMLV_stim_ slope-dependent effects in terms of dimensionality, decoding accuracy, and representational consistency within and across subjects. In sum, the geometry of trial-to-trial variability mediates the efficiency vs. robustness tradeoff in both monkey and mouse V1.

### Overdispersion from Poisson can enhance robustness

Neural spiking has long been compared to a Poisson process^2,4,36,51^, which has equal mean and variance. If all spike count distributions were Poisson, both LMLV_stim_ and LMLV_neuron_ would have a slope of 1 and a y-intercept of 0. So far, we have focused on the fact that the intrinsic geometry of neural variability had a characteristic LMLV_stim_ slope of 1 (Fig. 1). However, the LMLV_stim_ y-intercept was also highly consistent, with mean ± standard deviation of 0.14 ± 0.05 among natural scenes (Fig. 1a *right*). The fact that LMLV_stim_ y-intercept is greater than 0 is consistent with most neurons being overdispersed from Poisson (variance > mean). Accordingly, LMLV_neuron_ slopes were generally greater than 1 (Fig. 1a *left*; mean ± standard deviation 1.13 ± 0.21), and y-intercepts were generally greater than 0 (Fig. 1a *right*; mean ± standard deviation 0.17 ± 0.20).

To understand the impact of the overdispersion, we created pseudo-populations of Poisson neurons. We sampled spike counts from either the mean-matched or the variance-matched Poisson distributions. To have a control case that closely resembles the data, we sampled spike counts from the best-fit negative binomial distribution when variance > mean, or the best-fit Poisson distribution when variance ≤ mean (NB+P model). We used a Gaussian copula to incorporate pairwise noise correlations (see Methods). As most spike count distributions were overdispersed in the as-is data, there was a general downward shift onto the y = x diagonal of the LMLV_stim_ plot in the mean-matched Poisson model (decreased variance; Fig. 5a), and a general rightward shift onto the y = x diagonal in the variance-matched Poisson model (increased mean; Fig. 5a). As expected, the NB+P model behaved similarly to the as-is data (Fig. 5a-g).

**Fig. 5:**
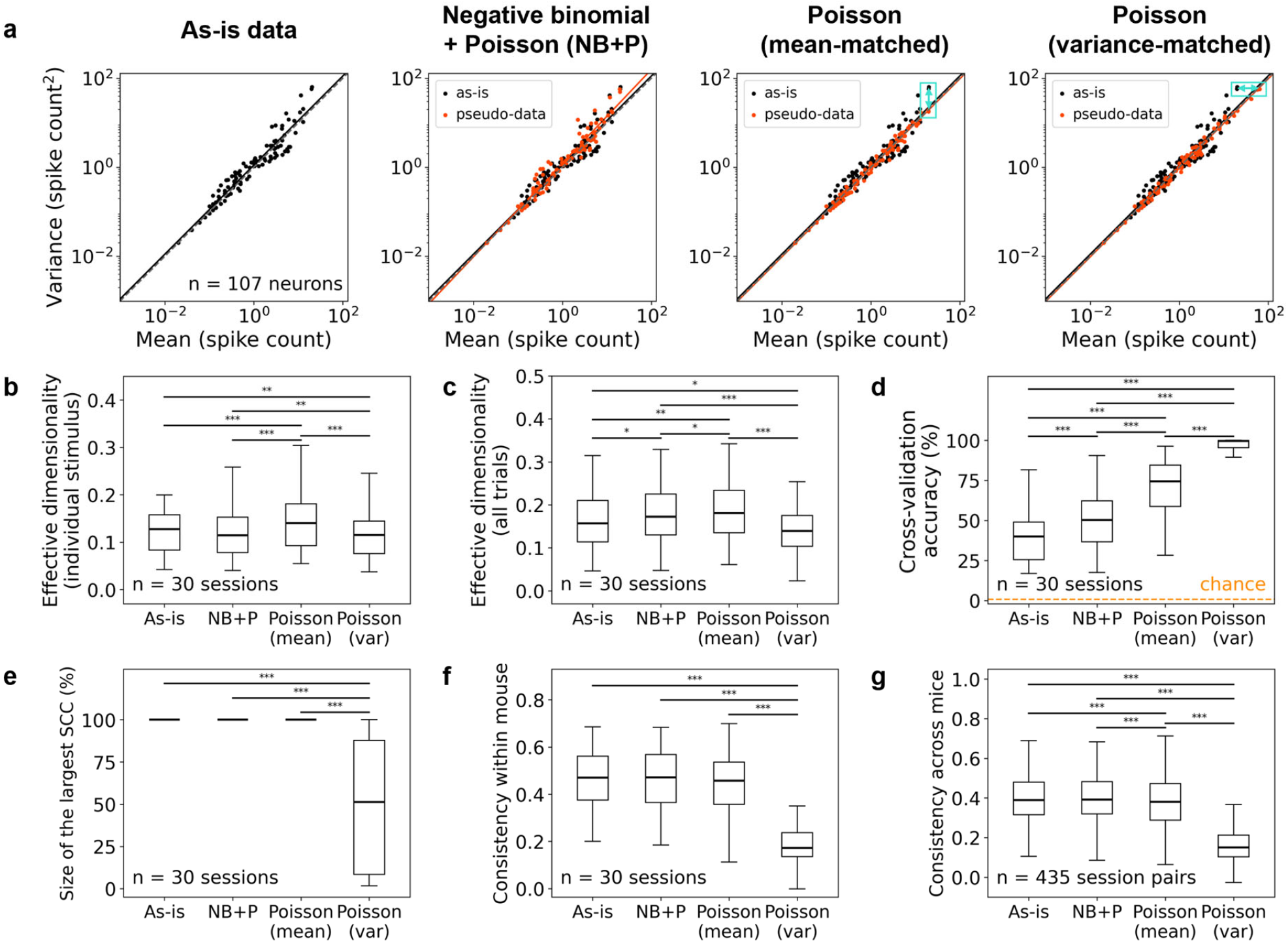
Overdispersion from Poisson can enhance robustness. **a** LMLV_stim_ scatterplots of an example natural scene image in an example session. The *black* dots denote as-is data. The *orange* dots denote the mean and variance of spike counts sampled from best-fit negative binomial or Poisson (‘NB+P’), mean-matched Poisson or variance-matched Poisson distributions. The *black* and *orange* lines are the lines fit by linear regression to the *black* and *orange* dots, respectively. The *turquoise* arrows indicate downward (‘mean-matched’) or rightward (‘variance-matched’) shift of dots. **b, c** Effective dimensionality of individual stimulus manifolds (**b**) and all stimulus manifolds (**c**). **d** 10-fold cross-validation classification accuracy of linear SVM decoder. *Orange* dotted line indicates chance level (1/119 ≈ 0.008). **e** Size of the largest SCC in the stimulus manifold graph. **f, g** Consistency of RSMs between distinct neuronal subsets within each mouse (**f**) and between pairs of mice (**g**). **b-g** Median and interquartile range (IQR) across sessions (**b-f**) or session pairs (**g**) were plotted. The asterisks (*) on top indicate significant differences (*p < 0.05, **p < 0.01, ***p < 0.001, two-sided Wilcoxon signed-rank test across sessions or session pairs, Holm-Bonferroni corrected).

The mean-matched Poisson model had smaller variances, leading to smaller and higher-dimensional stimulus manifolds (Fig. 5b, c). Consistent with these geometric changes, the decoding accuracy was higher (Fig. 5d). However, these geometric changes were not sufficient to create discontinuities in the visual representation space, as shown in the size of the largest SCC (Fig. 5e). Accordingly, representational consistency within a mouse was not significantly different from the as-is data (Fig. 5f). In contrast, representational consistency across mice was significantly lower in the mean-matched Poisson model compared to the as-is data (Fig. 5g). This suggests that overdispersion from Poisson contributes to consistent representations of visual similarity between different mice.

In the variance-matched Poisson model, the dimensionality slightly decreased (Fig. 5b, c), but the right-shift of spike count means led to better decoding accuracies (Fig. 5d). The shift of the stimulus manifold centroids away from the origin also created discontinuities in the representation space (Fig. 5e), resulting in less consistent visual representations both within and across mice (Fig. 5f, g).

In sum, spike count distributions were generally overdispersed from Poisson. Both mean-matched and variance-matched Poisson models have lower representational consistency across mice, and the variance-matched Poisson model additionally have lower representational consistency within a mouse. These results suggest that overdispersion from Poisson is beneficial for robust representation of visual similarity.

Next, we asked whether different neurons contribute differentially to the efficiency and robustness of visual representations. The fact that LMLV_stim_ has a consistent slope of 1 indicates that the neural population can be characterized by a single Fano factor. Yet, neurons had a wide range of Fano factors (Fig. 6a). Thus, we reasoned that neurons must balance each other to achieve a constant Fano factor at the population level. If so, we can break the balance by taking different subsets of neurons.

**Fig. 6:**
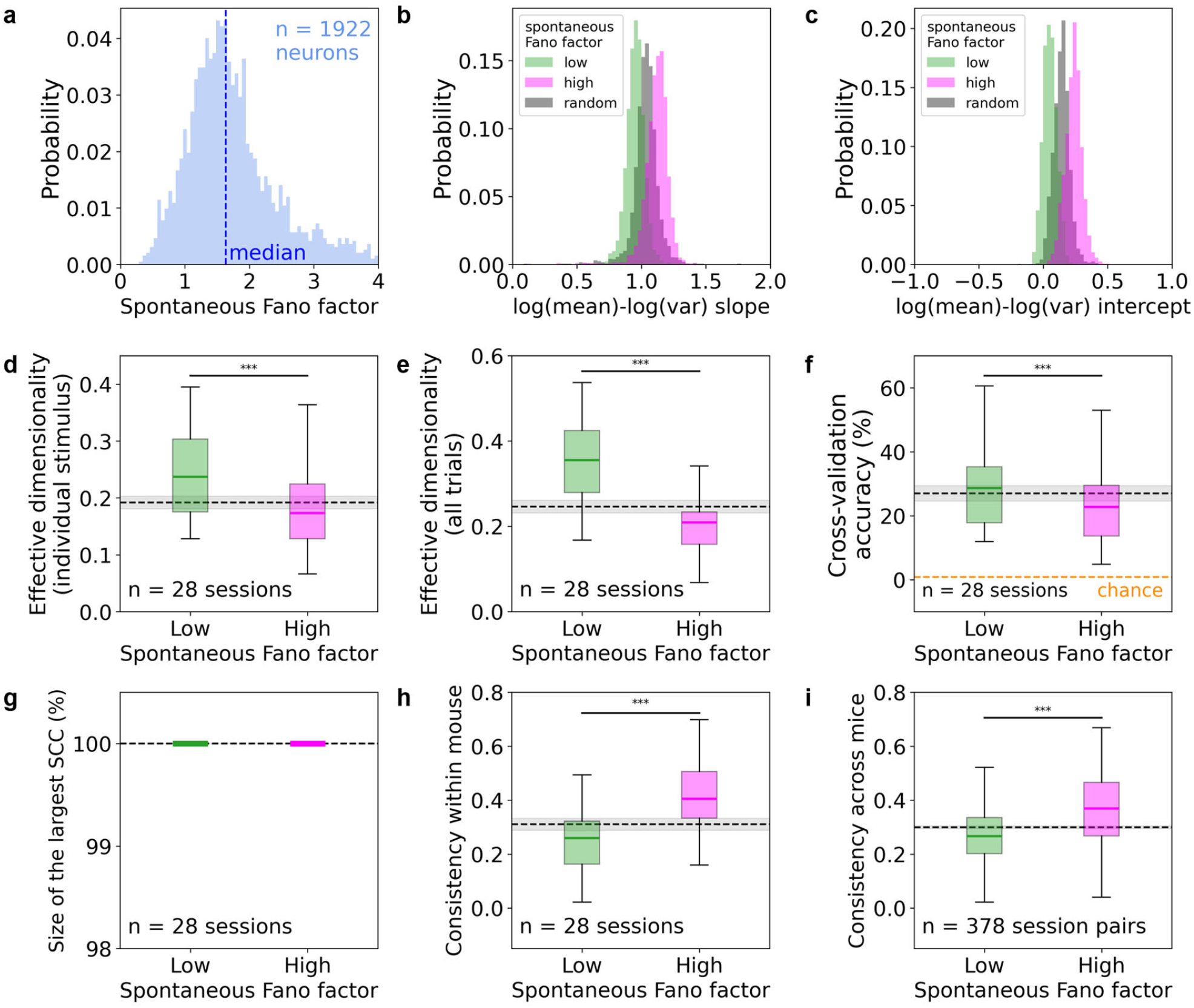
Overdispersed neurons are less efficient but more robust. **a** Histogram of Fano factors for V1 neurons pooled across 30 sessions of 30 mice. Fano factor of spike counts in 250 ms bins of prolonged spontaneous periods (≥ 5 min) was calculated for each neuron. **b, c** Histograms of LMLV_stim_ slopes (**b**) and y-intercepts (**c**) fit across V1 neuronal groups with different spontaneous Fano factors. Neurons in each session were divided in half based on spontaneous Fano factors (‘low’ in *green*, ‘high’ in *magenta*). The ‘random’ group consists of the same number of neurons randomly subsampled from each session (*gray*). Stimuli in 30 sessions of 30 mice were pooled. **d, e** Effective dimensionality of individual stimulus manifolds (**d**) and all stimulus manifolds (**e**). p = 1.02×10^-6^ (**d**) and p = 5.22×10^-8^ (**e**), two-sided Wilcoxon signed-rank test. **f** 10-fold cross-validation classification accuracy of linear SVM decoder. *Orange* dotted line indicates chance level (1/119 ≈ 0.008). p = 7.45×10^-9^, two-sided Wilcoxon signed-rank test. **g** Size of the largest SCC in the stimulus manifold graph. Sizes of the largest SCC were 100% across all sessions in both the ‘Low’ and ‘High’ groups. **h, i** Consistency of representational similarity matrices (RSMs) between distinct neuronal subsets within each mouse (**h**) and between pairs of mice (**i**). p = 5.53×10^-5^ (**h**) and p = 2.95×10^-32^ (**i**), two-sided Wilcoxon signed-rank test. **d-i** Median and IQR across sessions (**d-h**) or session pairs (**i**) were plotted. The asterisks (*) on top indicate significant differences (*p < 0.05, **p < 0.01, ***p < 0.001, two-sided Wilcoxon signed-rank test across sessions or session pairs). The *gray* dotted lines and shades indicate mean ± SEM across sessions or session pairs of the random group. Only sessions with 10 neurons in each group were analyzed (sessions with ≥ 20 neurons; 28 out of 30 sessions).

When we divided V1 neurons in half based on their Fano factor during prolonged spontaneous periods (≥ 5 minutes), the neuronal subsets showed different LMLV_stim_ slopes and y-intercepts (Fig. 6b, c). In other words, the geometry of trial-to-trial variability was different between neurons with low vs. high spontaneous Fano factor (mean ± standard deviation for lower half 1.15 ± 0.26, for upper half 1.97 ± 0.52). We compared these subsets with a random subset that has the same number of neurons, representing the balanced population (spontaneous Fano factor mean ± standard deviation 1.55 ± 0.62). The overdispersed neurons, i.e., the subpopulation with higher spontaneous Fano factors, had lower dimensionality and lower decoding accuracy but higher representational consistency within and across mice (Fig. 6d-i). Taken together, neurons with lower and higher spike count dispersion contribute to efficiency and robustness, respectively, thereby balancing this tradeoff.

### Efficiency-robustness tradeoff leads to changes in the geometry of trial-to-trial variability in a neural network simulation

What are the neural mechanisms that determine the geometry of trial-to-trial variability? The key to addressing this question is the finding that the mean vs. variance relationship is heterogeneous at the single-neuron level (LMLV_neuron_) but homogeneous at the population level (LMLV_stim_). This finding suggests that even though feedforward sensory inputs drive various subsets of neurons with heterogeneous LMLV_neuron_ relationships, LMLV_stim_ of a population remains constant. This implies that the geometry of trial-to-trial variability is shaped by a recurrent network mechanism rather than a feedforward mechanism. To investigate potential network mechanisms, we leveraged neural network simulations.

We adapted a computational model of leaky integrate-and-fire neurons with clustered connectivity, where excitatory and inhibitory neurons formed locally balanced clusters^27^ (Fig. 7a). This network has random connectivity with stronger within-cluster than between-cluster synaptic weights. This model reproduces realistic metastable attractor dynamics^52^ and stimulus-driven variability quenching^37^. In this model, each cluster acts as an attractor, such that neurons in the same cluster encode redundant information. Therefore, we predicted that increasing the cluster size, i.e., number of neurons per cluster, would partition a fixed-size network into fewer clusters with stronger attractor dynamics, increasing the representational robustness. Further, we hypothesized that changing representational efficiency and robustness in a network simulation would lead to corresponding changes in LMLV_stim_ slope. If true, this would establish a reciprocal relationship between the representational efficiency vs. robustness axis and LMLV_stim_ slope.

**Fig. 7:**
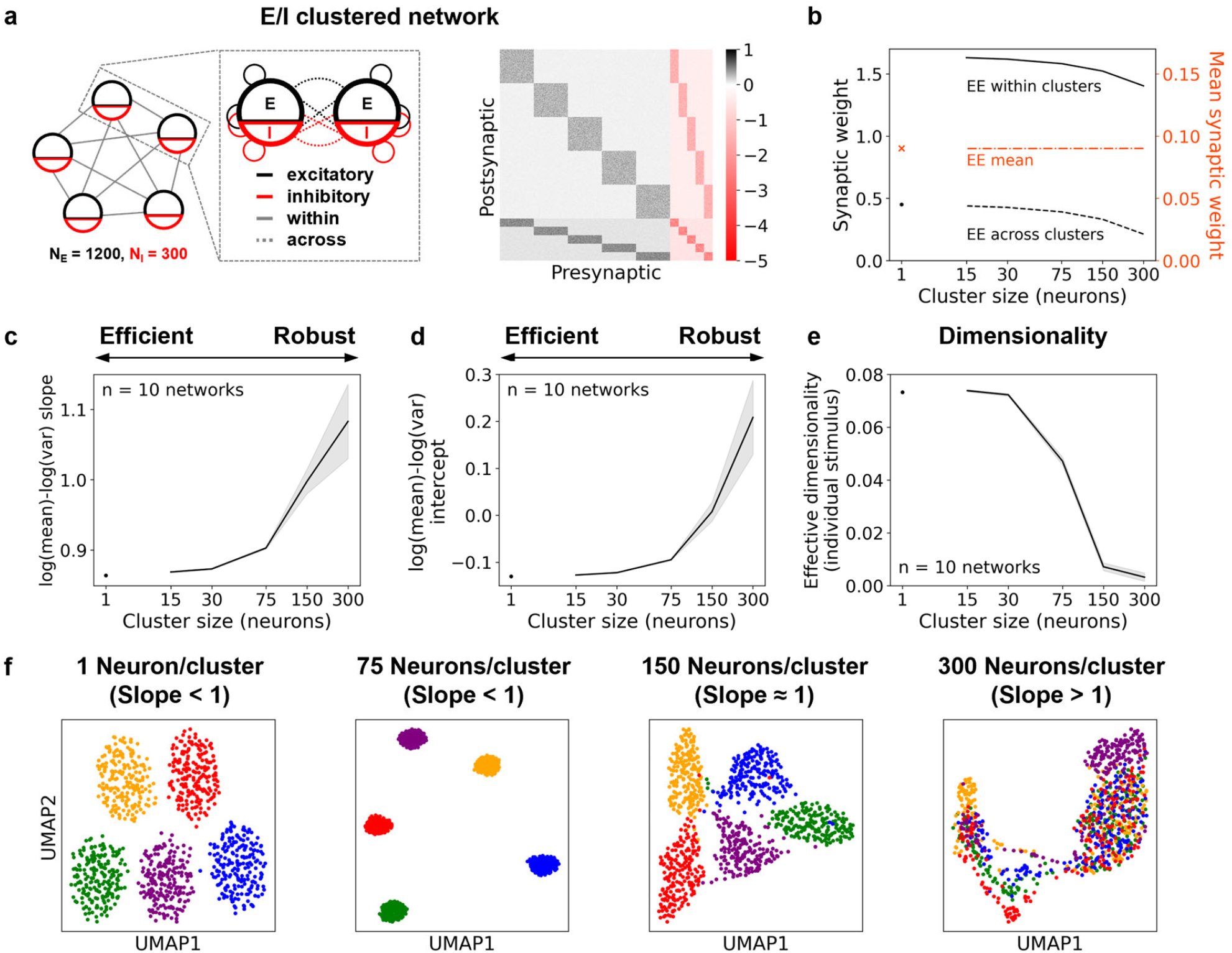
Efficiency-robustness tradeoff leads to changes in the geometry of trial-to-trial variability in a neural network simulation. **a** Schematic illustration of an example recurrent spiking neural network model with 5 clusters. *Left*: Both excitatory and inhibitory neurons were clustered by stronger synaptic weights, with fixed connection probability within and across clusters (p_EE_ = 0.2, p_EI_ = 0.5, p_IE_ = 0.5, p_II_ = 0.5). *Right*: Heatmap of synaptic weights in an example network with 5 clusters. *Black* and *red* indicates excitatory and inhibitory synaptic weights, respectively. **b** Within-cluster (left y-axis; *black*), across-cluster (left y-axis; *black*), and mean weights (right y-axis; *orange*) of excitatory-to-excitatory (‘EE’) synapses. For each cluster size, within- and across-cluster synaptic weights for connected pairs were set to maintain their difference and mean weight across all pairs, including unconnected pairs (zero weights). **c, d** LMLV_stim_ slope (**c**) and y-intercept (**d**) depending on cluster size. **e** Effective dimensionality of individual stimulus manifolds. **f** UMAP embedding of 5 stimulus manifolds in an example network. Each of the 5 colors is one stimulus manifold and each dot corresponds to a trial (200 trials per stimulus). **c-e** Mean ± SEM across networks was plotted as a function of the cluster size.

Each network comprised of 1,200 excitatory neurons and 300 inhibitory neurons. The cluster size was kept uniform within a network, with each cluster comprising of 80% excitatory neurons and 20% inhibitory neurons. The exception was when the number of neurons per cluster was 1, which is equivalent to a network with no clusters and Erdős–Rényi connectivity^53^. Across all networks, the mean synaptic weights were held constant, as well as the within-cluster vs. between-cluster weight differences (Fig. 7b). To examine stimulus-driven mean vs. variance relationships, we simulated five stimuli, with each stimulus driving a fifth of the clusters, corresponding to 240 excitatory neurons and 60 inhibitory neurons.

As we increased the cluster sizes, we observed an increase in LMLV_stim_ slopes and y-intercepts (Fig. 7c, d). Importantly, the geometric changes in trial-to-trial variability were consistent with those observed through LMLV_stim_ slope manipulations: As the cluster sizes increased, LMLV_stim_ slope increased, accompanied by decreasing dimensionality and increasing representational overlap (Fig. 7e, f). In addition, the decoding accuracy decreased as the cluster sizes increased, consistent with the accompanying increase in LMLV_stim_ slope (Supplementary Fig. 16e). Meanwhile, representational consistency between random subsets of neurons within a network tended to increase when the cluster sizes increased, plateauing near LMLV_stim_ slope 1 (Supplementary Fig. 16f). This is likely because increasing cluster sizes results in fewer clusters with stronger attractor dynamics, leading to greater robustness within a network. In contrast, representational consistency across networks tended to decrease with increasing number of neurons per cluster (Supplementary Fig. 16g). This is likely because when cluster sizes are larger, each stimulus recruits fewer clusters, and random differences in attractors cannot be effectively averaged out. Therefore, RSMs become less uniform (Supplementary Fig. 16c), and representational consistency across networks decreases.

In sum, parametric changes in the efficiency vs. robustness axis in a simulated network led to corresponding changes in the geometry of trial-to-trial variability, establishing a reciprocal relationship. These results further suggest that in a cortical network with recurrently connected neural ensembles, the granularity of neural ensembles may be a key determinant of the geometry of trial-to-trial variability.

## Discussion

Neural variability obeys a simple rule: The mean and the variance of spike counts in response to repeated presentations of the same sensory stimulus are linearly related in the log-log scale across a neural population, with a slope of 1. We investigated the functional implication of this highly consistent log-mean vs. log-variance relationship across neurons per stimulus (LMLV_stim_). To this end, we systematically varied the LMLV_stim_ slope across simultaneously recorded V1 neurons and examined the geometry of the visual representation space in V1. When the slope increased, the dimensionality of the neural state space decreased. Further, representations of different images became increasingly overlapping with increasing slope, impairing discriminability. At slopes below 1, the representation space was discontinuous, precluding a robust neural code. At slope 1, discriminability was maximal within the constraint of global continuity, which was achieved by allocating more variance towards dissimilar representations and less variance towards similar representations. Finally, the visual representation space in V1 was most consistent at slope 1, both within and across mice. Together, our findings indicate that even though the intrinsic geometry of trial-to-trial variability is suboptimal for efficient coding, it is optimal for robust coding.

We found that distinct subsets of simultaneously recorded neurons within one brain area encode consistent sensory information. This coding consistency peaked when the LMLV_stim_ slope was around 1. How does the brain benefit from such coding consistency between neuronal subsets within a brain region? A brain region typically communicates with several downstream areas. For example, mouse V1 sends monosynaptic projections to many downstream regions, including LM, AL, RL, AM, and PM^54^. In many cases, distinct downstream areas are targeted by distinct neurons^55,56^. For effective global information processing, it would be ideal to relay a unified message to all downstream targets.

We also found that the intrinsic geometry of trial-to-trial variability allows different individuals to have consistent representations of visual similarity. Prior literature has demonstrated representational consistency between individuals^57,58^, but these studies mostly focused on trial-averaged responses (so-called neural ‘signal’). In comparison, we focused on the role of trial-to-trial variability (neural ‘noise’) in representational consistency. Our result suggests that the internal models of the external sensory world can be consistent, even if sensory representations are mostly learned through individual experiences. Shared internal models could be a consequence of representational smoothness, ensuring that similar stimuli are represented similarly. In sum, our results imply that the intrinsic geometry of trial-to-trial variability may enable individual brains to build consistent representations of the objective reality despite individual differences in sensory experiences. Such consensus on the objective reality may be critical for building collective knowledge amongst groups of individuals^59^.

What is the neural mechanism underlying the incredibly consistent geometry of trial-to-trial variability? Our network simulation results suggest that in a network of clustered excitatory and inhibitory connectivity, the cluster size determines the LMLV_stim_ slope, as well as the balance between efficient vs. robust visual representations. In support of the model premise, the like-to-like connectivity rule predicts clustered connectivity in visual cortex, with stronger connections between neurons that have similar visual response profiles^60-62^. Further, this network model recapitulates metastable attractor dynamics observed in neural recordings^27^. As the ensemble size grows, the attractor dynamics gain strength, leading to more robust but less efficient representations, accompanied by an increase in LMLV_stim_ slope. The stability of the geometry of trial-to-trial variability suggests that there may be homeostatic cortical mechanisms that govern the clustered connectivity of neural ensembles.

How does variability change with learning or with familiarization to the sensory stimulus? A recent study found that stimulus manifolds shrink in all directions as the number of examples available for generalization increases^63^. We showed that the geometry of trial-to-trial variability does not change with familiarization (Fig. 1k-n). Together, these findings suggest that stimulus manifolds shrink in all directions proportionally, accompanied by a mean shift toward the origin of the neural state space. This isomorphic shrinkage may allow the representation of familiar stimuli to gain efficiency while maintaining robustness. More generally, these results suggest that the intrinsic geometry of trial-to-trial variability not only enables few-shot generalization, but also helps maintain the stability of representations during familiarization.

Many studies have investigated the influence of trial-to-trial variability on sensory representation and the neural code. Most of these studies focused on pairwise noise correlations^12,18,64,65^. Here, we employed a residual rescaling method that leaves pairwise noise correlations intact. When we destroyed pairwise noise correlations by shuffling the trial order of each stimulus for each neuron, the LMLV_stim_ slope-dependent trends remained largely consistent. These results demonstrate that pairwise noise correlations are only a part of whole story; it is important to consider the neural population geometry when assessing the role of trial-to-trial variability in the neural code^43,66^.

Our findings are in line with the idea that neural variability is a feature of optimal perceptual inference^16,26,67,68^. In the sampling-based Bayesian framework, stimulus manifolds in the visual representation space correspond to the posterior distributions in perceptual space^23,67,69,70^. Differentiable posteriors are desired for probabilistic Bayesian inference, since small differences in sensory input should not lead to large perceptual differences. Thus, visual representation space must be smooth for probabilistic inference. While stimulus manifolds must overlap for visual representations to be smooth, overlapping stimulus manifolds also imply poor discriminability. Therefore, the sampling-based Bayesian framework predicts that continuous stimulus manifolds with minimal overlap are ideal for probabilistic inference, suggesting that the intrinsic geometry of trial-to-trial variability is well suited for this computation. Together, these results indicate that the geometry of trial-to-trial variability plays a key role in sensory encoding under the framework of probabilistic perceptual inference.

## Methods

### Datasets and visual stimuli

#### Extracellular electrophysiology recordings of mouse visual cortex during passive viewing

We analyzed the Allen Brain Observatory Visual Coding Neuropixels dataset^28^, where extracellular electrophysiology recordings were made from awake head-fixed mice that were passively viewing a battery of visual stimuli (https://portal.brain-map.org/circuits-behavior/visual-coding-neuropixels). Six Neuropixels^71^ probes were targeted to retinotopically aligned regions in six visual cortices, including primary visual cortex (V1) and 5 higher visual areas (lateromedial LM, rostrolateral RL, anterolateral AL, posteromedial PM, and anteromedial AM). Details of the experimental procedure and data processing can be found in ^28^ and the online white paper (https://brainmapportal-live-4cc80a57cd6e400d854-f7fdcae.divio-media.net/filer_public/80/75/8075a100-ca64-429a-b39a-569121b612b2/neuropixels_visual_coding_-_white_paper_v10.pdf).

Among single units, only those that passed the quality control criteria were analyzed (‘isi_violations’ < 0.5, ‘amplitude_cutoff’ < 0.1, and ‘presence_ratio’ > 0.9)^28^. We analyzed only V1 units except for the natural scenes data, for which we examined all 6 visual cortical areas. We excluded invalid sessions for each area and stimulus type, which had stimulus failures, data loss for one probe, or gain fluctuations, as described in the online white paper. We also excluded sessions that have no units in the analyzed area.

We analyzed the spike counts during the stimulus presentation period (0–250 ms after stimulus onset) of the ‘natural scenes’ and ‘static gratings’ blocks from the ‘Brain Observatory 1.1’ sessions (32 sessions, each session corresponds to a mouse). In addition, we analyzed prolonged spontaneous periods where a gray screen was presented for ≥ 5 minutes.

In the ‘natural scenes’ block, 118 different natural scene images and 1 gray-screen ‘blank’ image were presented for 250 ms, in a randomized order with no gray screen period between trials. All 119 stimuli were presented 50 times, except for one session where the stimulus repeats ranged from 47 to 50. For this session, we randomly sampled the minimum number of trials (47 trials) from each stimulus before conducting the decoding analysis and the representational similarity analysis (RSA). We focused on the natural scenes data unless otherwise noted.

In the ‘static gratings’ block, 120 different static gratings and 1 gray-screen ‘blank’ image were presented for 250 ms, in a randomized order with no gray screen period between trials. Static gratings were shown at 6 orientations (0°, 30°, 60°, 90°, 120°, 150°), 5 spatial frequencies (0.02, 0.04, 0.08, 0.16, 0.32 cycles/degree), and 4 phases (0, 0.25, 0.5, 0.75). Each of 120 static gratings were presented 39–50 times, while the blank image was presented 180–197 times.

We analyzed the ‘drifting gratings’ and ‘natural movie’ blocks from the ‘Functional Connectivity’ sessions (26 sessions, each session corresponds to a mouse). In addition, we analyzed a long spontaneous block where a gray screen was presented for 30 minutes.

In the ‘drifting gratings’ block, 8 different drifting gratings were presented for 2 s, in a randomized order with 1 s gray screen period between trials. Drifting gratings were shown at 4 directions (0°, 45°, 90°, 135°) and 2 contrasts (0.1, 0.8), with a temporal frequency of 2 Hz. All 8 stimuli were presented 75 times. Spike counts during 0–250 ms after stimulus onset were used. In the ‘natural movie’ block, we analyzed ‘natural movie one’ presentations, which refers to a 30 s movie clip taken from the opening scene of the movie Touch of Evil^72^, repeated 60 times.

#### Extracellular electrophysiology recordings of monkey V1 during passive viewing

We analyzed extracellular electrophysiology recordings from 2 awake head-fixed monkeys that were passively viewing natural scenes (32 sessions, 15 and 17 sessions from monkey M1 and M2, respectively)^30^. In this dataset, acute recordings of V1 and V4 were conducted using 32-channel linear silicon probes (NeuroNexus V1x32-Edge-10mm-60-177). Details of the experimental procedure and data processing can be found in ^30^.

We analyzed V1 single unit spike counts during 40–160 ms after stimulus onset to account for the spike latency, as provided in the publicly available dataset. A total of 24,075 natural scenes were shown: The ‘train-set’ and ‘validation-set’ contained a total of 24,000 images, each presented at most once; the ‘test-set’ contained 75 images, each repeated 17–65 times. Because our analysis required repeated presentations, we analyzed only the ‘test-set’ trials. Note, Cadena et al.^30^ defined a ‘trial’ as a 2.4 s period where 15 images was sequentially shown for 120 ms with no gray screen period between images, flanked by 300 ms of gray screen at the beginning and end. The ‘trial’ was successful if the monkey maintained fixation throughout the 2.4 s period. We analyzed these successful fixation ‘trials’. For consistency with other datasets, we treated each repeat of a ‘test-set’ image as a trial.

The number of V1 single units recorded in each session was much fewer than the mouse datasets (238 neurons across 15 sessions in M1, 220 neurons across 17 sessions in M2). In Fig. 1, we pooled all neurons in each monkey. In Supplementary Fig. 15, we pooled and subsampled neurons in each monkey to create pseudo-sessions of 110 neurons (30 pseudo-sessions, 15 per monkey). We randomly bootstrapped 50 trials for each neuron and stimulus, which increased the trial count and removed noise correlations altogether.

#### Extracellular electrophysiology recordings of mouse V1 during change detection task while viewing familiar and novel images

We analyzed the Allen Brain Observatory Visual Behavior Neuropixels dataset^32^, where extracellular electrophysiology recordings were made from awake head-fixed mice that were conducting a natural scenes change detection task (https://brain-map.org/our-research/circuits-behavior/visual-behavior). Six Neuropixels^71^ probes were targeted to retinotopically aligned regions in six visual cortices (V1, LM, RL, AL, PM, and AM), similar to the Allen Brain Observatory Visual Coding Neuropixels dataset^28^. Details of the experimental procedure and data processing can be found in ^32^ and the online white paper (https://cdn.prod.website-files.com/689cfbd308fa7373b604d290/691cd5932e9982f63e2beaf6_visualbehaviorneuropixels_technicalwhitepaper.pdf).

Mice were trained to perform a natural scene change detection task, in which licking in response to a change in image identity was rewarded with water. Each image was presented for 250 ms with 500 ms gray screen between images, repeated 322–933 times with ∼3% probability of omission (gray screen). Image changes occurred randomly after 4 or more repetitions, leading to adaptation of neural activity in trials following the change trial. Each recording session consisted of an active task block followed by a passive replay block. We analyzed both active and passive viewing conditions in ‘familiar’ and ‘novel’ sessions. In ‘familiar’ sessions, the image set contained 8 familiar natural scene images used throughout training (Fig. 1l, n). In ‘novel’ sessions, the image set contained 6 novel and 2 familiar natural scene images. We only analyzed the novel images in ‘novel’ sessions (Fig. 1k, m). We analyzed V1 single unit spike counts during the 250 ms visual presentation window. We excluded two sessions with no V1 units, resulting in a total of 101 sessions from 54 mice.

#### Two-photon calcium imaging of mouse V1 during passive viewing

We analyzed the calcium fluorescence and deconvolved events in Allen Brain Observatory Visual Coding Optical Physiology dataset^73,74^. Two-photon calcium imaging was performed at 30 Hz frame rate, in six visual cortical areas (V1, LM, RL, AL, PM, and AM) of multiple transgenic mouse lines, while mice were passively viewing similar visual stimuli with the Allen Brain Observatory Visual Coding Neuropixels dataset^28^. Details of the experimental procedure and data processing can be found in ^73^ and the online white paper (https://s3.amazonaws.com/webflow-prod-assets/689cfbd308fa7373b604d290/68ee796f6f627db2434e459c_Documentation_Brain_Observatory-Visual_Coding_Overview.pdf).

We analyzed V1 excitatory neurons in 7 Emx1-IRES-Cre;Camk2a-tTA;Ai93(TITL-GCaMP6f) mice (1,894 neurons in 10 sessions) and 15 Slc17a7-IRES2-Cre;Camk2a-tTA;Ai93(TITL-GCaMP6f) mice (3,082 neurons in 17 sessions). We focused on the ‘natural scenes’ block where the same natural scene images as the Allen Brain Observatory Visual Coding Neuropixels dataset were presented; 118 natural scenes and 1 gray image were shown for 250 ms in random order with no intervening period between images, each repeated 50 times. dF/F for each trial of a stimulus was computed using the average fluorescence of the previous 1s as the baseline F_0_, and we used the average dF/F across frames within each trial. We also used deconvolved ‘event’ counts^40^ for each trial.

### Linear regression of log-mean vs. log-variance of spike counts

For each stimulus and session, we calculated the across-trial mean and variance of spike counts. To examine log-mean vs. log-variance relationship across neurons per stimulus (LMLV_stim_), we fit the slope and intercept by ordinary least squares (OLS) linear regression of log-mean and log-variance across neurons for each stimulus (Fig. 1). Conversely, to examine the log-mean vs. log-variance relationship across stimuli per neuron (LMLV_neuron_), we fit linear regression of log-mean and log-variance across stimuli for each neuron (Fig. 1). We excluded neuron-stimulus combinations with zero spike counts across all trials.

To test whether the LMLV_stim_ and LMLV_neuron_ relationships are reproduced in two-photon calcium imaging data, we fit linear regression to average dF/F and deconvolved event counts^40^ for natural scenes. We excluded neuron-stimulus combinations with mean dF/F ≤ 0, or zero event counts across trials.

### Comparison of Modulated Poisson and Log-log-linear model for log-mean vs. log-variance relationship across stimuli per neuron

We fit the Modulated Poisson model to the mouse natural scenes data (Supplementary Fig. 1)^75^. Modulated Poisson model is a doubly stochastic process model, in which the spike count (*k*) of neuron *n* for stimulus *s* follows a Poisson distribution while the mean firing rate (*μ*_*n,s*_) emerges as the product of stimulus-driven feedforward drive (*f*_*n*_(*s*)) and stimulus-independent multiplicative gain (*G*_*n*_):

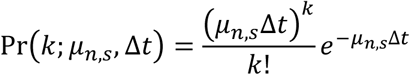

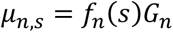

Δ*t* indicates the time window of spike count. *G*_*n*_ is assumed to have an average of 1. Marginalizing over *G*_*n*_ results in a negative binomial distribution of the spike count (*Eq*. 1)^75^:

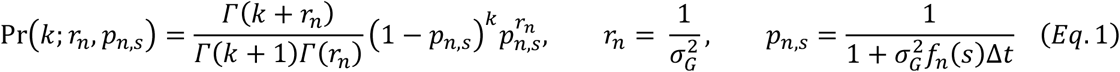

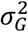 indicates the variance of gain *G*_*n*_ . The mean and variance of spike count follows a quadratic relationship across stimuli per neuron^75^:

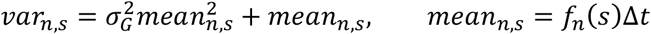

First, to test whether the natural scenes data follows the assumption that the multiplicative gain is independent of the stimulus, we fit *r*_*n,s*_ and *p*_*n,s*_ to each neuron-stimulus combination by maximum likelihood estimation (MLE). We then estimated the gain variance 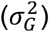 as the reciprocal of fitted *r*_*n,s*_ (*Eq*. 1) and compared the standard deviation and median of 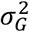 across stimuli per neuron vs. across neurons per stimulus (Supplementary Fig. 1b, c).

Next, we fit the Modulated Poisson model (‘MP’) and the log-log-linear model (‘LLL’) to each neuron in the mouse natural scenes data (Supplementary Fig. 1e, f). In the MP model for neuron *n*, one *r*_*n*_ and a set of *p*_*n,s*_ values for all stimuli (*s*) were jointly fit; specifically, we identified the *r*_*n*_ and *p*_*n,s*_ values that minimized the across-stimulus sum of the log-likelihood of spike count distributions. Neurons that do not follow the supralinearity of MP model, i.e., neurons for which the variance across all trials of all stimuli is lower than the mean, were excluded.

The LLL model assumed that the mean vs. variance relationship for neuron *n* and stimulus *s* follows log *var*_*n,s*_ = *a*_*n*_ log *mean*_*n,s*_ + *b*_*n*_, and that, for each stimulus, the spike counts follow a negative binomial distribution if *var*_*n,s*_ > *mean*_*n,s*_ and a Poisson distribution if *var*_*n,s*_ ≤ *mean*_*n,s*_. The parameters of the negative binomial and Poisson distributions were estimated by the method of moments, and the values of *a*_*n*_ and *b*_*n*_ were determined by minimizing the across-stimuli sum of the log-likelihood of spike count distributions.

We compared the two models by 10-fold cross-validation of the negative log-likelihood (NLL), Akaike information criterion (AIC), and root mean squared error (RMSE) of spike count variance (Supplementary Fig. 1f). For each neuron, model parameters were fit using 90% of the natural scene trials (train set) and then tested either on the remaining 10% of the natural scene trials (test set) or on all static grating trials recorded in the same session; the latter was intended to assess generalizability across stimulus types. Because the number of test trials differed between these two conditions, we averaged across trials within each stimulus when calculating the NLL and AIC. The number of parameters (*m*) in AIC was calculated as the sum of the number of distribution parameters (*m*_*d*_) and the number of stimuli (*m*_*s*_) when tested within natural scenes; *m*_*d*_ was not included when tested using static grating trials because the distribution parameters fit to natural scene trials were used.

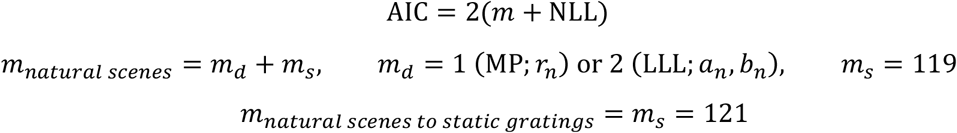

### Manipulation of slope by residual rescaling

While preserving the mean spike count of each neuron and the total log-variance (across-neuron sum of log-variance per stimulus), we manipulated the stimulus-specific spike count variances to change the LMLV_stim_ slope from 0 to 2 (incrementing by 0.1). To make such pseudo-data, we rescaled the residual of each trial (spike count minus the across-trial mean) by multiplying it with an appropriate constant. The constant was determined as follows.

Let *a*_*s*_ and *b*_*s*_ be the best-fit slope and intercept for a specific stimulus *s*, respectively. Given a new target slope 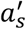, new variance 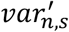 for neuron *n* and intercept 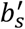 must satisfy the following equation:

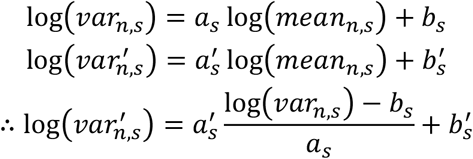

We can determine a single 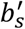 for each target slope 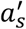 that preserves the total log-variance as follows:

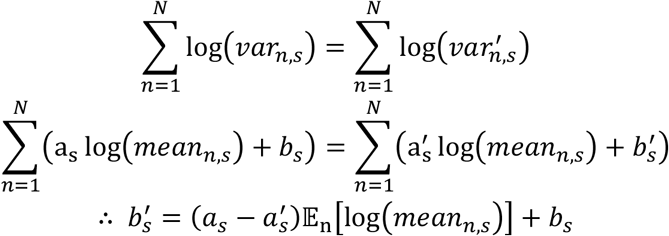

*N* denotes the number of neurons for each session. For each stimulus, we determined 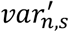 for neuron *n* and multiplied each trial’s residual by 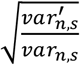 to make the spike count variance equal to 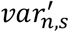 .

### Visualization of stimulus manifold geometry

Trial-by-trial responses to natural scenes for all simultaneously recorded neurons can be represented in the neural state space. Each axis in the neural state space corresponds to a neuron, and each point corresponds to the neural activity vector on a single trial (#neurons-by-1 vector of spike counts). The stimulus manifold refers to the collection of points in the neural state space for all repetitions of a given stimulus. UMAP^41^, a nonlinear dimensionality reduction technique, was used to visualize the geometric changes of the stimulus manifolds in the neural state space following changes in the LMLV_stim_ slope (Fig. 2e, 7g, Supplementary Fig. 4). We used Python package ‘umap’ for implementation of UMAP.

In Fig. 2e, we plotted all 50 repetitions of 2 example natural scenes in an example session. To get a clear intuition for overlap between stimulus manifolds, we added density contour of each stimulus manifold using kernel density estimation (KDE)^76,77^. We used Python function ‘seaborn.kdeplot’ for plotting of KDE contours^78^. In Fig. 7f, we plotted all 200 repetitions of 5 stimuli in an example network. In Supplementary Fig. 4, we treated each 400 ms bin of the natural movie as a single stimulus and plotted all trials of all stimuli in an example session. After slope manipulation, we compared the UMAP embedding of the as-is (no slope manipulation) data with that of slope 0, 1, and 2.

### Effective dimensionality

The effective dimensionality was defined as follows:

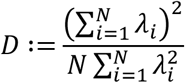

*N* and *λ*_*i*_ refers to the number of neurons and the *i*th eigenvalue of the covariance matrix, respectively. If we remove *N* from the denominator of this expression, it becomes the expression derived in ^79^, referred to as the ‘participation ratio’^80,81^. We normalized by dividing by *N* to control for the different number of neurons in each session. Effective dimensionality has a maximum value of 1, which is reached when explained variances are the same across all principal components.

### Overlap percentage between stimulus manifolds

We quantified the overlap percentage between individual stimulus manifolds, adapting the metric from ^42^ (Fig. 3a). Specifically, for each point in stimulus manifold A, we drew a hypersphere with a radius equal to the bottom 5th percentile of pairwise distances among points in A. The union of these hyperspheres defined the volume of stimulus manifold A. Accordingly, the overlap percentage of stimulus manifold B relative to A was calculated as the fraction of points in B that fell within the volume of A. Note that the overlap percentage is asymmetrical; the overlap percentage of stimulus manifold B relative to A is different from that of stimulus manifold A relative to B.

### Decoding of visual stimuli

To assess the neural discriminability of visual stimuli, we leveraged linear support vector machine (SVM) for decoding stimulus identity. We used Python class ‘sklearn.svm.SVC’^82^ for implementation. Within each session, trials were divided into training and test sets for stratified 10-fold cross-validation. In each of the 10 iterations, neural activity of each neuron in the training and test sets were centered by the average neural activity across training trials. Classification accuracy was computed as the proportion of correctly classified test trials, and the final performance was reported as the cross-validation average accuracy.

In Supplementary Fig. 6a *right*, we trained the decoder with 90% of the trials in slope 0 data and tested with the remaining 10% of every slope data including slope 0.

### Strongly connected component (SCC) of stimulus manifold network

To assess the continuity of the visual representation space, we defined a directed graph of stimulus manifolds, where each stimulus manifold was treated as a node, and edges between nodes were determined based on the presence of overlap between stimulus manifolds. For example, for stimuli A and B, if the overlap percentage of stimulus A relative to stimulus B is greater than zero, we defined a directed edge from node B to node A in the graph. For each session, we calculated the size of the largest SCC^83^. The SCC was defined as the subgraph in which every node could reach every other node via a directed path.

### Orthogonal variance

We adapted the method from ^45^ to measure the variance of a stimulus manifold in the direction orthogonal to the hyperplane linearly separating it against another manifold (Supplementary Fig. 5a-d).

Orthogonal vector **r**_**orth**_ of a point (trial) in the stimulus A manifold relative to the stimulus B manifold was defined as follows:

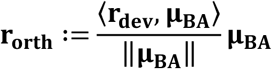

Here, **μ**_**BA**_ refers to **μ**_**B**_ **− μ**_**A**_, where **μ**_**A**_ and **μ**_**B**_ are the centroid vectors of the two manifolds. **r**_**dev**_ : = **r**_**A**_ **− μ**_**A**_ refers to the deviation vector of the neural activity vector **r**_**A**_ for a stimulus A trial. ⟨·,·⟩ indicates the dot product of two vectors, and ‖·‖ denotes the *L*^2^ norm of a vector.

Orthogonal distance was defined as the *L*^2^ norm of **r**_**orth**_, i.e., the magnitude of the deviation vector component that is parallel to the vector between two centroids of a stimulus manifold pair. Orthogonal variance of stimulus A relative to stimulus B was calculated as the variance of orthogonal distances, normalized by the sum of variances across neurons.

To measure the inter-centroid geodesic distances, the centroid vectors of all stimulus manifolds were concatenated to the neural activity matrix (#neurons-by-#trials) and the geodesic distances were calculated using the scikit-learn Isomap class (‘sklearn.manifold.Isomap’). Then, across all stimulus pairs, we calculated the Spearman correlation between the inter-centroid geodesic distance and the orthogonal variance (Supplementary Fig. 5b-d).

### Manifold alignment

To probe the alignment of dominant directions of each stimulus manifold and its nearest neighbor stimulus (local alignment), we calculated the absolute value of the cosine similarity between PC1’s of each stimulus manifold and its neighbor stimulus manifold (Supplementary Fig. 5e). The nearest neighbor stimulus was defined based on the geodesic distance between the centroids of the stimulus manifolds.

To probe the alignment of dominant directions of each stimulus manifold and the global manifold (global alignment), we calculated the absolute value of the cosine similarity between PC1’s of each stimulus manifold and the global manifold excluding that stimulus (Supplementary Fig. 5f).

### Overlap consistency

To examine how consistent the similarity relationship of stimulus manifolds are between distinct neuronal subsets within and across mice, we calculated the consistency of overlap percentages. For comparison of neuronal subsets within a mouse, we randomly divided the neurons of each session into two subsets of equal size. For each subset, we computed an overlap percentage matrix sized #stimuli-by-#stimuli. Then, we calculated the Spearman correlation of the two matrices. We repeated the random partitioning 10 times and averaged the 10 Spearman correlations within each session. For pairs of mice, we computed the overlap percentage matrix for each session and calculated the Spearman correlation of the matrices between all session pairs.

We also performed a leave-one-mouse-out (LOMO) analysis to quantify overlap consistency between different mice (Supplementary Fig. 6b *left*). We computed the Spearman correlation between each session’s overlap percentage matrix and the average overlap percentage matrix of the remaining sessions.

### Representational similarity analysis (RSA)

We performed RSA^48^ to see how the change in the geometry of trial-to-trial variability affects the robustness of representational similarity structure between different neuronal subsets, both within and across brains (Fig. 4b). We created a representational similarity matrix (RSM) for each neuronal subset and measured the consistency of RSMs between two distinct subsets within each session, or between session pairs, using the Spearman correlation. For within-mouse analysis, we repeated the random partitioning of neurons 10 times and averaged the 10 Spearman correlations within each session.

The process for creating an RSM for each neuronal subset was as follows. First, we constructed a 3-dimensional the neural activity matrix sized #neurons-by-#stimuli-by-#repetitions. We randomized the order in the trial repetition dimension of the neural activity matrix. Then, the trial repetition dimension was indexed from 1 through *r*, where *r* is the number of trial repeats (50 in most sessions). Next, for every possible trial index pair, we calculated the cosine similarity between neural activity vectors of size *N* (number of neurons) for all stimulus pairs, creating a similarity matrix sized #stimuli-by-#stimuli. For example, if the trial index pair is (10, 33), a similarity matrix was created by calculating the cosine similarity between the 10th trial vector and the 33rd trial vector for all stimulus pairs. Given *r* repetitions, the total number of trial index pairs is 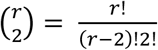, so total 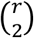 matrices were created and averaged to acquire a single RSM for each neuronal subset.

To quantify representational consistency between different mice, we also performed a leave-one-mouse-out (LOMO) analysis (Supplementary Fig. 6b *right*). We computed the Spearman correlation between each session’s RSM and the average RSM of the remaining sessions.

To investigate representational consistency between distinct visual cortical areas, we conducted RSA between all pairs of 6 visual cortical areas (V1, LM, RL, AL, PM, and AM; Supplementary Fig. 13). For within-mouse comparison, we randomly sampled *N* neurons from each of the two areas, where *N* is equal to half the number of neurons in the area with fewer neurons. We repeated this procedure 10 times for each area pair and averaged the 10 Spearman correlations of RSMs within each session. For comparison of mouse pairs, we computed the Spearman correlations of RSMs between different areas of different mice. When comparing different LMLV_stim_ slopes, we manipulated the slope based on the as-is slopes fit within each area in each session.

For drifting gratings, we compared the results of RSA using RSMs with and without diagonal elements (i.e., representational consistency within each stimulus), both within and across mice (Supplementary Fig. 14d, e, g, h).

### Exclusion and partitioning of neurons based on receptive field locations

We re-evaluated the LMLV_stim_ and LMLV_neuron_ relationships and LMLV_stim_ slope manipulation analyses after excluding neurons whose receptive fields are located outside the monitor (Supplementary Fig. 9a-f). For each V1 neuron in ‘Brain Observatory 1.1’ sessions, the receptive field was estimated by fitting a two-dimensional Gaussian distribution to the 9 × 9 spatial map of spike counts evoked by Gabor stimuli presented at the 81 locations^28^. Neurons were classified as on-screen if the center of the fitted receptive field fell within the screen boundaries. Applying this criterion left 46.3 ± 20.1 on-screen neurons (mean ± standard deviation across sessions), corresponding to 70.7 ± 16.8% of all V1 neurons. We restricted the analyses to sessions containing at least 10 neurons.

Furthermore, to quantify the representational consistency between neuronal subsets with non-overlapping receptive fields, we divided neurons in each session into left vs. right or top vs. bottom halves (Supplementary Fig. 9g-l). As the monitor spanned the receptive field center’s azimuth from 10° to 90° and elevation from -30° to 50° (Supplementary Fig. 9g, j), we performed RSA between neuronal subsets divided based on azimuth of 50° or elevation of 10°, both within and across mice. We restricted the analyses to sessions with at least 10 neurons in both halves.

### Manipulation of LMLV_stim_ slope by sampling spike counts from discrete probability distributions

Manipulation of LMLV_stim_ slope by residual rescaling did not guarantee biologically plausible (i.e., non-negative integer) spike counts. Therefore, to replicate the effects of residual rescaling with non-negative integer spike counts, we randomly sampled spike counts from discrete probability distributions (Supplementary Fig. 11). Depending on the target mean and variance of each neuron-stimulus combination that achieves the target slope, we sampled 50 spike counts from negative binomial if variance ≥ mean × 1.05, Poisson if mean × 0.95 < var < mean × 1.05, or binomial if var ≤ mean × 0.95. Spike counts were sampled independently for each neuron and stimulus, ignoring pairwise noise correlations.

### Sampling spike counts from Poisson or negative binomial distribution fit to as-is data

To investigate the influence of overdispersion of spike counts from Poisson distribution, we sampled spike counts from Poisson or negative binomial distributions (Fig. 5). For each neuron-stimulus combination, the mean of the Poisson distribution was matched to the mean of the spike count distribution in the mean-matched Poisson model, and the variances were matched in the variance-matched Poisson model. To make a spike count sampling model that follows the as-is data more closely, we fit a negative binomial distribution to a neuron-stimulus combination if the spike count variance > mean, and a Poisson distribution if variance ≤ mean, by maximum likelihood estimation (MLE; ‘NB+P’).

To attempt to preserve pairwise noise correlations, we used Gaussian copula when sampling from the fitted distributions^84,85^. First, we performed Cholesky decomposition of the Pearson noise correlation matrix **R**_**s**_ for each stimulus *s* of the as-is data.

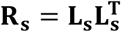

We then imposed this empirical correlation structure on random samples **Z**_**s**_, sized #neurons-by-#samples, drawn from an independent multivariate standard normal distribution. The number of samples was matched to the number of trials in the as-is data (i.e., 50 samples).

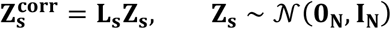

Here, *N* denotes the number of neurons in each session, **0**_**N**_ and **I**_**N**_ denote an *N* - dimensional zero vector and an *N*-by-*N* identity matrix, respectively. We next applied the marginal cumulative distribution function (*ϕ*) of standard normal distribution to each neuron *n* in the correlated Gaussian samples 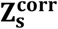.

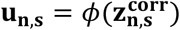

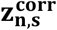 indicates the neuron *n*’s gaussian samples for stimulus *s*, i.e., *n*th row vector of 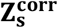. Finally, we applied the marginal inverse cumulative distribution function 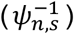 of either negative binomial or Poisson distribution to each neuron *n* and stimulus *s* to obtain the final spike count vector (**y**_**n**,**s**_; *Eq*. 2). This procedure allowed each neuron to have the desired marginal distribution.

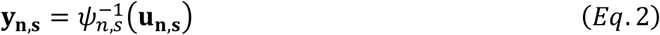

For mean-matched and variance-matched Poisson data, 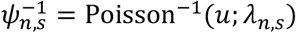 where *λ*_*n,s*_ is the parameter of Poisson probability mass function for spike count *k*:

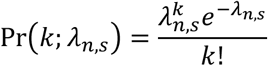

*λ*_*n,s*_ is equal to mean spike count *mean*_*n,s*_ (mean-matched) or variance of spike count *var*_*n,s*_ (variance-matched). For NB+P data we used the parameters fit by MLE:

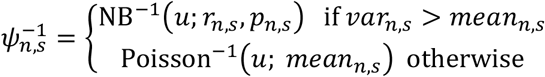

*r*_*n,s*_ and *p*_*n,s*_ are the parameters of negative binomial probability mass function for spike count *k*:

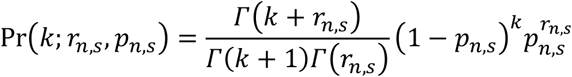

Note that this sampling method slightly underestimated pairwise noise correlations because spike counts were obtained by discretizing continuous samples from the Gaussian copula (*Eq*. 2; mean ± standard deviation across 30 sessions 0.10 ± 0.04 for as-is; 0.08 ± 0.03 for NB+P; 0.08 ± 0.03 for mean-matched Poisson; 0.09 ± 0.04 for variance-matched Poisson)^85^. Total log-variance was also not entirely preserved (mean ± standard deviation across 30 sessions 8.28 ± 6.25 for as-is; 8.52 ± 6.04 for NB+P; -1.46 ± 6.99 for mean-matched Poisson; 7.78 ± 6.06 for variance-matched Poisson). Slight differences in NB+P pseudo-data and as-is data were likely due to slight differences in pairwise noise correlations and total log-variance (Fig. 5c, d).

### Partitioning and comparison of neurons based on spontaneous Fano factor

For each spontaneous period (≥ 5 minutes) in the Brain Observatory 1.1 sessions, 250 ms spike counts were calculated and all such periods were concatenated to a #neurons-by-#bins matrix. We then calculated, for each neuron, the Fano factor of spike counts across all time bins and split neurons within each session into lower and higher half. As a control, we generated a ‘random’ group by randomly subsampling the same number of neurons within each session (Fig. 6).

### Simulation of clustered recurrent spiking neural network

We adapted a recurrent spiking neural network model introduced in ^86^, achieving realistic E-I balance, metastable attractor dynamics and stimulus-evoked variability quenching (Fig. 7, Supplementary Fig. 16). The network consists of excitatory and inhibitory leaky integrate-and-fire (LIF) neurons. Each neuron’s subthreshold membrane potential *V* evolved according to the following equation:

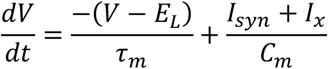

Here, *E, τ*_*m*_, *I*_*syn*_, *I*_*x*_, and *C*_*m*_ denote the resting potential (*E*_*L*_), membrane time constant (*τ*_*m*_ ), recurrent synaptic input current (*I*_*syn*_), external input current (*I*_*x*_), and membrane capacitance (*C*_*m*_) (Table 1). Note that *I*_*x*_ was set as a constant current, rather than a Poisson input, to ensure that all neural variability arises from inside the recurrent network. *I*_*x*_ was determined by multiplying a constant by *I*_*th*_, the current needed to reach the threshold *V*_*th*_ in the absence of synaptic input (Table 1):

**Table 1:** Network parameters used in simulation.

| Parameter | Unit | Value |
| --- | --- | --- |
| $N$ | - | 1,200 ($N_E$), 300 ($N_I$) |
| $E_L$ | mV | 0 |
| $V_{th}$ | mV | 15 |
| $V_R$ | mV | 0 |
| $C_m$ | pF | 1 |
| $\tau_m$ | ms | 20 ($E$), 10 ($I$) |
| $\tau_{syn}$ | ms | 3 ($E$), 2 ($I$) |
| $\tau_r$ | ms | 5 |
| $p_{EE}$ | - | 0.2 |
| $p_{EI}, p_{IE}, p_{II}$ | - | 0.5 |
| $g$ | - | 1.2 |
| $PSP_{max}^{EE}$ | mV/pA | 2.15 |
| $PSP_{max}^{EI}$ | mV/pA | 1.55 |
| $PSP_{max}^{IE}$ | mV/pA | 1.79 |
| $PSP_{max}^{II}$ | mV/pA | 1.34 |
| $I_x$ | pA | $1.25I_{th}$ ($E$), $0.78I_{th}$ ($I$) |
| $Q_{base}$ | - | 6 |
| $J_{E,base}^+$ | - | 3.2 |
| $R_j$ | - | 0.75 |

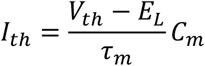

When *V* reached the threshold *V*_*th*_, a spike was emitted, and *V* was clamped to the reset potential *V*_*r*_ for an absolute refractory period *τ*_*r*_ . The synaptic current *I*_*syn*_ received by neuron *i* evolved according to the following equation:

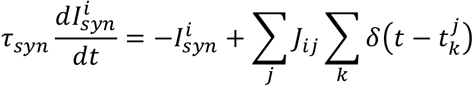

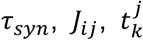 and *δ* denote the synaptic time constant (*τ*_*syn*_), synaptic weight from neuron *j* to *i* (*J*_*ij*_), time of the *k*th spike of presynaptic neuron 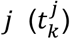, and Dirac delta function (*δ*).

Each network consisted of *N*_*E*_ = 1,200 excitatory neurons and *N*_*I*_ = 300 inhibitory neurons, connected randomly according to fixed connection probabilities (*p*_*EE*_ = 0.2, *p*_*EI*_ = *p*_*IE*_ = *p*_*II*_ = 0.5). The synaptic weights were determined such that first, 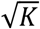 excitatory spikes during a short time, where *K* denotes the average number of connections a neuron receives, are sufficient to elicit a postsynaptic spike^87^; and second, the excitatory and inhibitory inputs to each neuron are balanced^88^.

In the absence of clustering (i.e., when each neuron formed its own cluster), the scale-free synaptic weights were defined for each of the four synapse types (*Eq*.3 **−** 6).

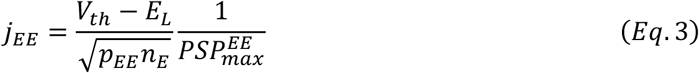

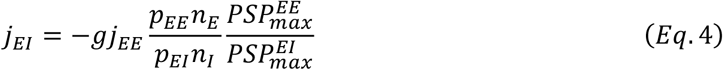

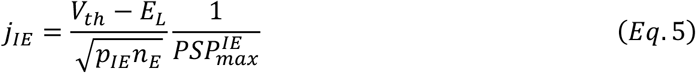

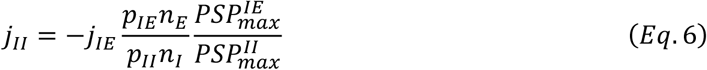

Here, *g* and *PSP*_*max*_ denote the rela
tive strength of inhibition (*g*) and peak postsynaptic potential for a synaptic weight of 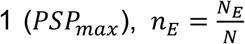 and 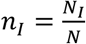. For each synapse type, *PSP*_*max*_ depends only on *τ*_*m*_ and *τ*_*syn*_ ^86^. Because these two parameters were kept fixed across all networks in our analyses, *PSP*_*max*_ was also constant for all networks (Table 1). The final weights *J*_*EE*_, *J*_*EI*_, *J*_*IE*_, and *J*_*II*_ were obtained by dividing *j*_*EE*_, *j*_*EI*_, *j*_*IE*_, and *j* by 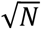, where *N* = *N*_*E*_ + *N*_*I*_ is the total number of neurons in the network.

In clustered networks, excitatory and inhibitory neurons within the same cluster had stronger synaptic weights than those in different clusters. Specifically, the synaptic weights were obtained by multiplying the weight values of the non-clustered network by the clustering factors 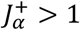 and 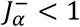, for within- and across-cluster synapses, respectively 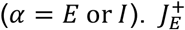 and 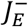 were applied to EE synapses, whereas 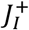 and 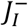 were applied to the other three synapse types. 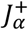 and 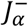 were constrained to keep the mean synaptic weight constant (*Eq*. 7; Fig. 7b):

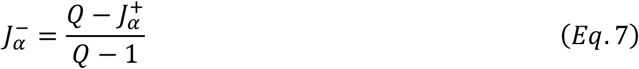

*Q* denotes the number of clusters. To examine how changing the coding efficiency and robustness affects the LMLV_stim_ slope, we varied Q with fixed *N* while maintaining the difference between 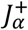 and 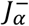 constant (*Eq*. 8; Fig. 7b):

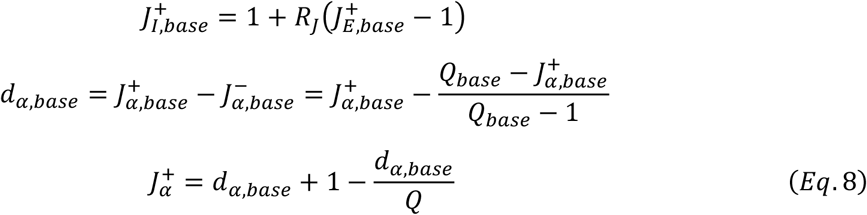

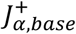 and 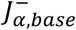 denote the default values of 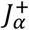 and 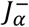 used in ^86^ to match experimental data. *R*_*J*_ is a constant smaller than 1 applied to make realistic firing rates^86^. Combined with *Eq*.7, *Eq*.8 ensures that the difference of 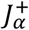 and 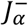 always matches *d*_*α,base*_, i.e., the difference of 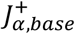 and 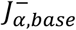 . All the parameter values except 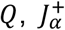 and 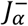 were kept the same as those used in ^86^ (Table 1).

For each of Q = 5, 10, 20, 50, and 100, we simulated networks with 10 different random seeds. For a given Q, the 10 networks had identical within- and across-cluster weights for all four synapse types and differed only in their connection patterns. To simulate 5 stimuli, we divided the excitatory neurons into 5 non-overlapping groups (i.e., neurons belonging to Q/5 clusters), and increased the external input *I*_*x*_ by 0.1 for 250 ms in one group at a time. Each of the 5 stimuli were repeated 200 times, for a total of 1000 trials, in a randomized order with zero inter-stimulus interval. For RSA within and across networks, we constructed a RSM for each neuronal subset by averaging 10,000 cosine similarity matrices of trial-to-trial spike count vectors (Supplementary Fig. 16c, f, g).

## Data and code availability

Data analyzed in this study are publicly available (see Methods)^29-31,38^.

Code is available at: https://github.com/jhk4839/Neural_variability_robustness

## Acknowledgments

We thank Dr. Kenneth Harris for helpful discussions. This work was supported by the Samsung Science and Technology Foundation (SSTF-BA2302-07), and the National Research Foundation of Korea (NRF) grant funded by the Korean government (MSIT) (RS-2024-00358070, RS-2024-00413689, RS-2023-00301976).

---

**Supplementary Fig. 1:**
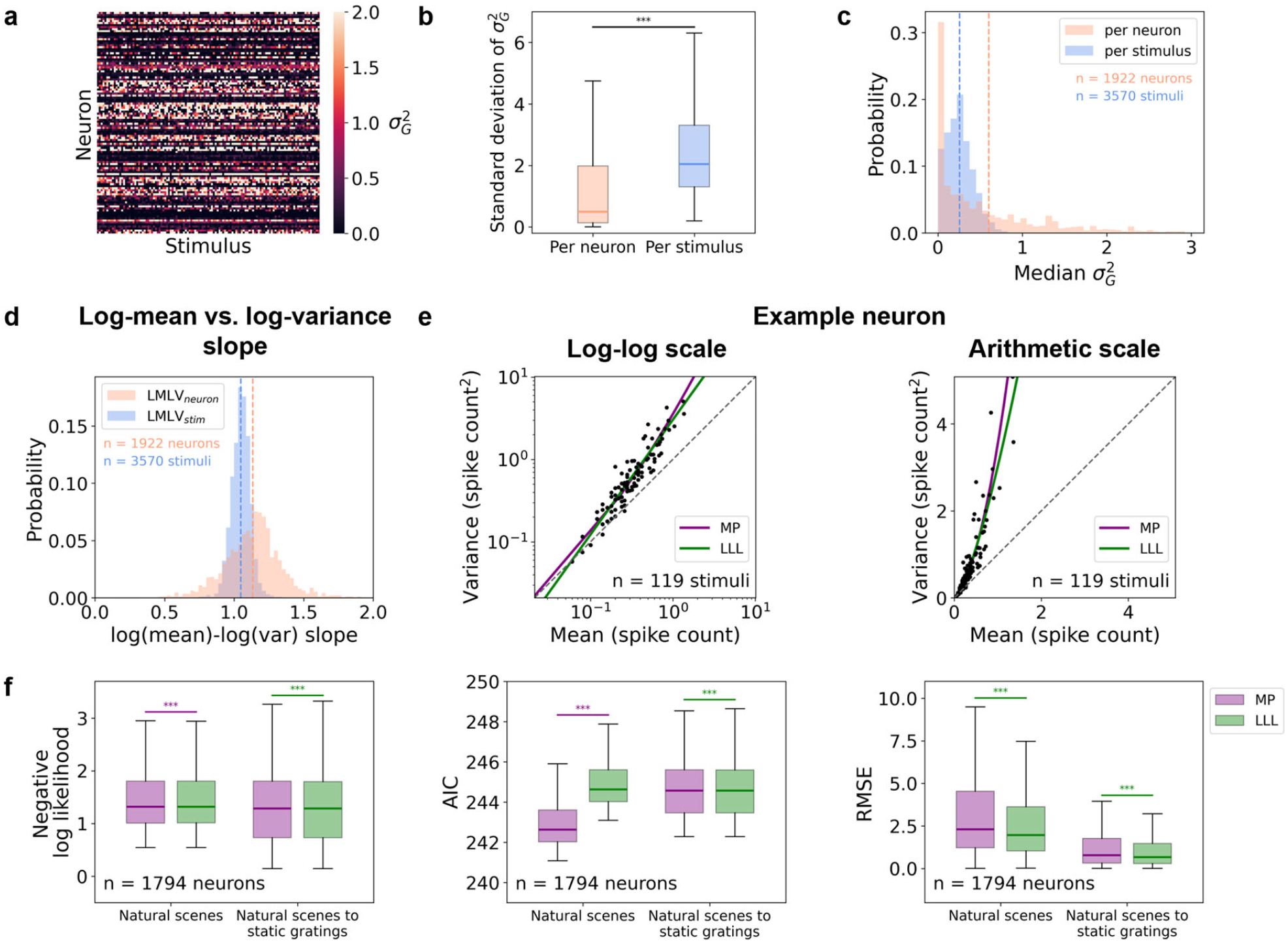
Log-mean vs. log-variance relationship across stimuli per neuron (LMLV_neuron_) is heterogeneous, and is consistent with the Modulated Poisson model. **a** Heatmap of gain variance 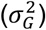 in an example session. A negative binomial NB(*r, p*) was fit to each V1 neuron and stimulus by maximum likelihood estimation, and gain variance was calculated as the inverse of 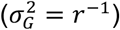. **b** Box plots of standard deviation of gain variance. Standard deviation was computed across stimuli per neuron (*pink*) or across neurons per stimulus (*blue*). Median and IQR were plotted. p = 3.85×10^-207^, two-sided Wilcoxon rank-sum test. **c** Histograms of median gain variance. Median gain variance was computed across stimuli per neuron (*pink*) or across neurons per stimulus (*blue*). **d** Histograms of log-mean vs. log-variance slopes across neurons per stimulus (LMLV_stim_; *blue*) and across stimuli per neuron (LMLV_neuron_; *pink*) for mouse V1 responses to natural scenes. Same as **Fig. 1a** *left*. **b-d** V1 neurons and stimuli in in 30 sessions of 30 mice were pooled. In **c** and **d**, the dotted lines indicate the mean of each condition. **e** Mean vs. variance scatterplots of an example neuron in log-log (*left*) and arithmetic (*right*) scale. The *purple* (MP; Modulated Poisson) and *green* (LLL; log-log linear) lines indicate maximum likelihood estimation fits to the *black* dots. **f** Box plots of negative log-likelihood (*left*), Akaike information criterion (AIC; *center*), and root mean squared error of variance (RMSE; *right*) of Modulated Poisson (MP) and log-log linear (LLL) models. Each model was fit to a neuron with spike counts for natural scenes and tested using either held-out trials for natural scenes (‘Natural scenes’), or static gratings (‘Natural scenes to static gratings’), using 10-fold cross-validation. Median and IQR across V1 neurons pooled across 30 sessions were plotted, excluding neurons with lower variance than mean across all natural scene trials (83 out of 1,877 neurons, ∼4%). Two-sided Wilcoxon signed-rank test was applied to compare MP and LLL across neurons (n = 1,794): p = 1.02×10^-15^ for ‘Natural scenes’ and p = 2.91×10^-51^ for ‘Natural scenes to static gratings’ (negative log-likelihood), p = 1.21×10^-294^ for ‘Natural scenes’ and p = 2.91×10^-51^ for ‘Natural scenes to static gratings’ (AIC), p = 2.15×10^-158^ for ‘Natural scenes’ and p = 5.45×10^-52^ for ‘Natural scenes to static gratings’ (RMSE). Colors of asterisks indicate the better model (i.e., lower value). **b, f** The asterisks (*) on top indicate significant differences (*p < 0.05, **p < 0.01, ***p < 0.001).

**Supplementary Fig. 2:**
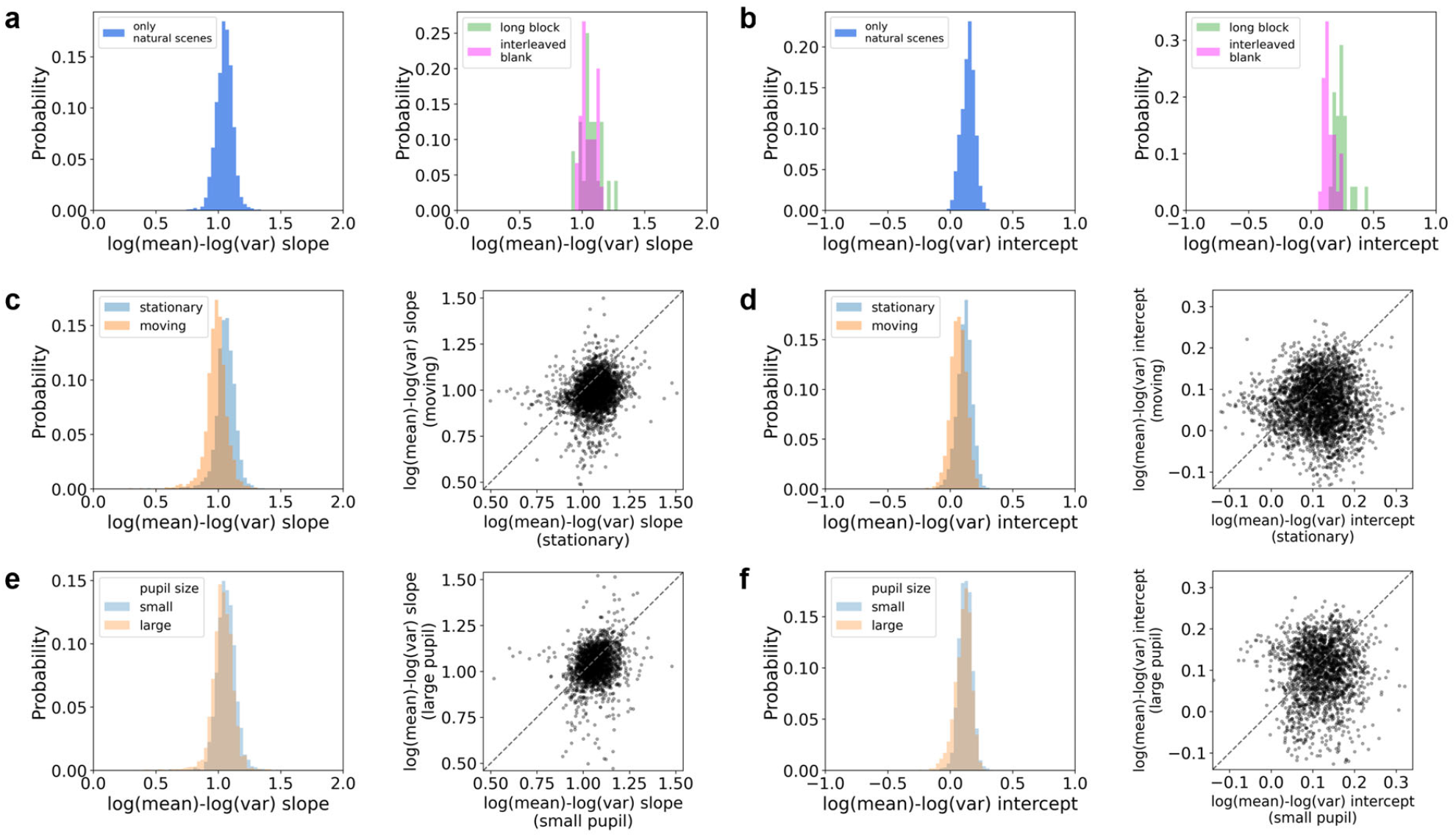
Log-mean vs. log-variance of spike counts across neurons depending on stimulus presence, locomotion, and pupil size. **a, b** Histograms of LMLV_stim_ slopes (**a**) and y-intercepts (**b**) across V1 neurons for natural scenes (*left*; 118 natural scenes) and gray screen periods (*right*). For gray screen periods, prolonged spontaneous block (30 min divided into non-overlapping 250 ms bins and 50 bins subsampled; ‘long block’ in *green*) and gray images interleaved with natural scenes (‘interleaved blank’ in *magenta*) were shown. Slopes were not significantly different across the 3 conditions (p = 0.23 for natural scenes vs. long block, p = 0.74 for natural scenes vs. interleaved blank, and p = 0.43 for long block vs. interleaved blank, two-sided Wilcoxon rank-sum test; **a**). Y-intercepts were significantly different between the long spontaneous block and either the natural scenes or interleaved blank trials (p = 6.52×10^-13^ for natural scenes vs. long block, p = 0.98 for natural scenes vs. interleaved blank, and p = 2.03×10^-7^ for long block vs. interleaved blank, two-sided Wilcoxon rank-sum test; **b**). **c, d** Histograms (*left*) and a scatterplot (*right*) of LMLV_stim_ slopes (**c**) and y-intercepts (**d**) across V1 neurons depending on mouse locomotion. Each trial was categorized as ‘moving’ if the average mouse speed was ≥ 1 cm/s, and ‘stationary’ otherwise. p = 4.20×10^-269^ (**c**) and p = 1.97×10^-163^ (**d**), two-sided Wilcoxon signed-rank test. **e, f** Histograms (*left*) and a scatterplot (*right*) of LMLV_stim_ slopes (**e**) and y-intercepts (**f**) across V1 neurons depending on mouse pupil size. Each trial was categorized as ‘small’ if the average mouse pupil size was smaller than the median value of all mice, and ‘large’ otherwise. p = 6.79×10^-40^ (**e**) and p = 4.19×10^-17^ (**f**), two-sided Wilcoxon rank-sum test. **a-f** Slopes or y-intercepts in all sessions were pooled (24 sessions of 24 mice for **a, b** ‘long ‘block’; 30 sessions of 30 mice for the rest).

**Supplementary Fig. 3:**
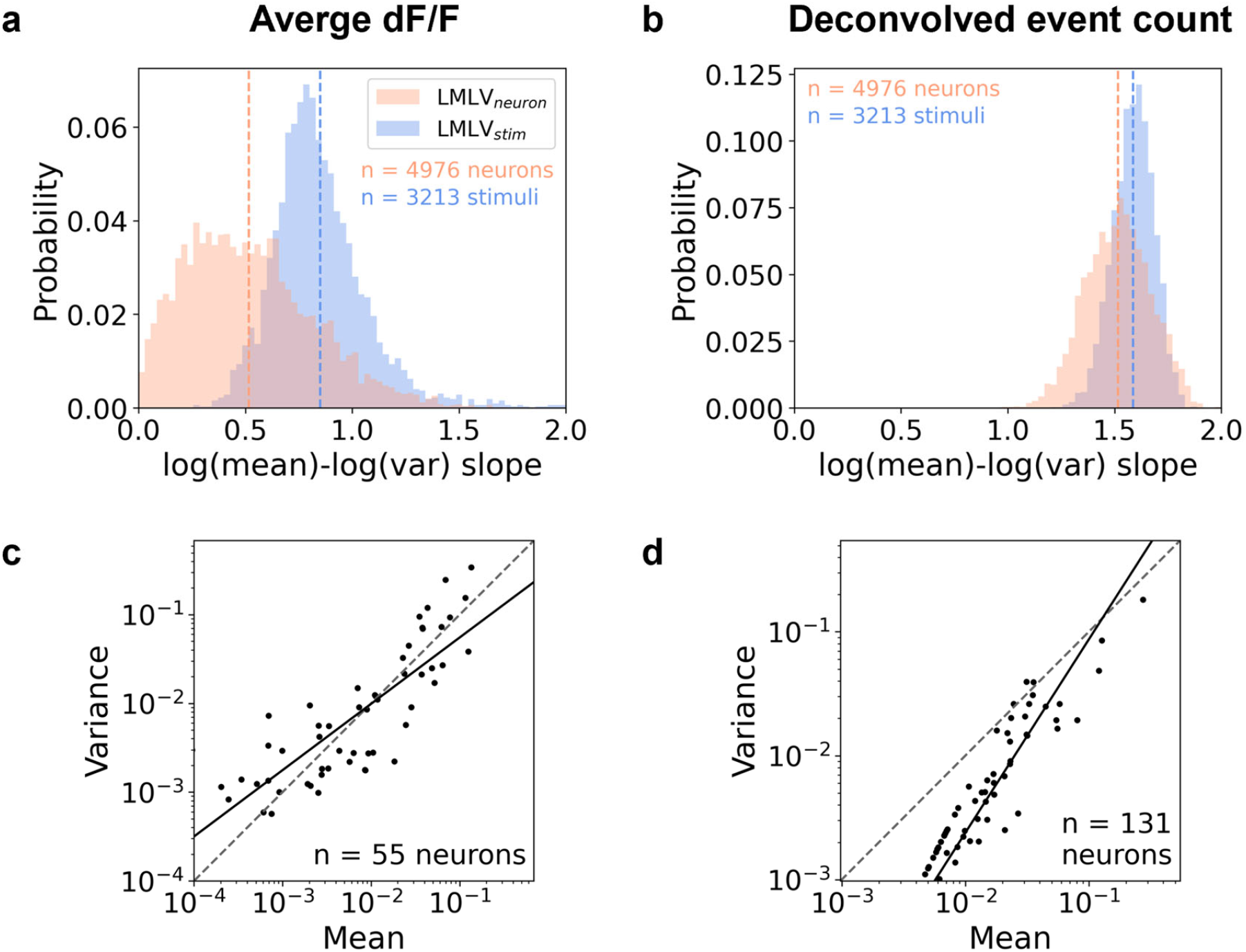
Mean vs. variance relationship of spike counts is not reproduced with two-photon calcium imaging. **a, b** Histograms of LMLV_stim_ and LMLV_neuron_ slopes across V1 neurons for average dF/F (**a**) and deconvolved event count (**b**) during natural scene presentations. 119 images (118 natural scenes and 1 gray image) were presented for 250 ms, each repeated 50 times. dF/F and deconvolved event were averaged and summed, respectively, during the stimulus presentation period. Stimuli and neurons in 27 sessions of 22 mice were pooled (10 sessions from 7 Emx1-IRES-Cre;Camk2a-tTA;Ai93(TITL-GCaMP6f) mice; 17 sessions from 15 Slc17a7-IRES2-Cre;Camk2a-tTA;Ai93(TITL-GCaMP6f) mice). The dotted lines indicate the mean of each condition. **c, d** LMLV_stim_ scatterplots of an example session and a natural scene, for average dF/F (**c**) and deconvolved event count (**d**). The black line is the line fit by linear regression of the black dots. Only the neurons with means higher than 0 were included.

**Supplementary Fig. 4:**
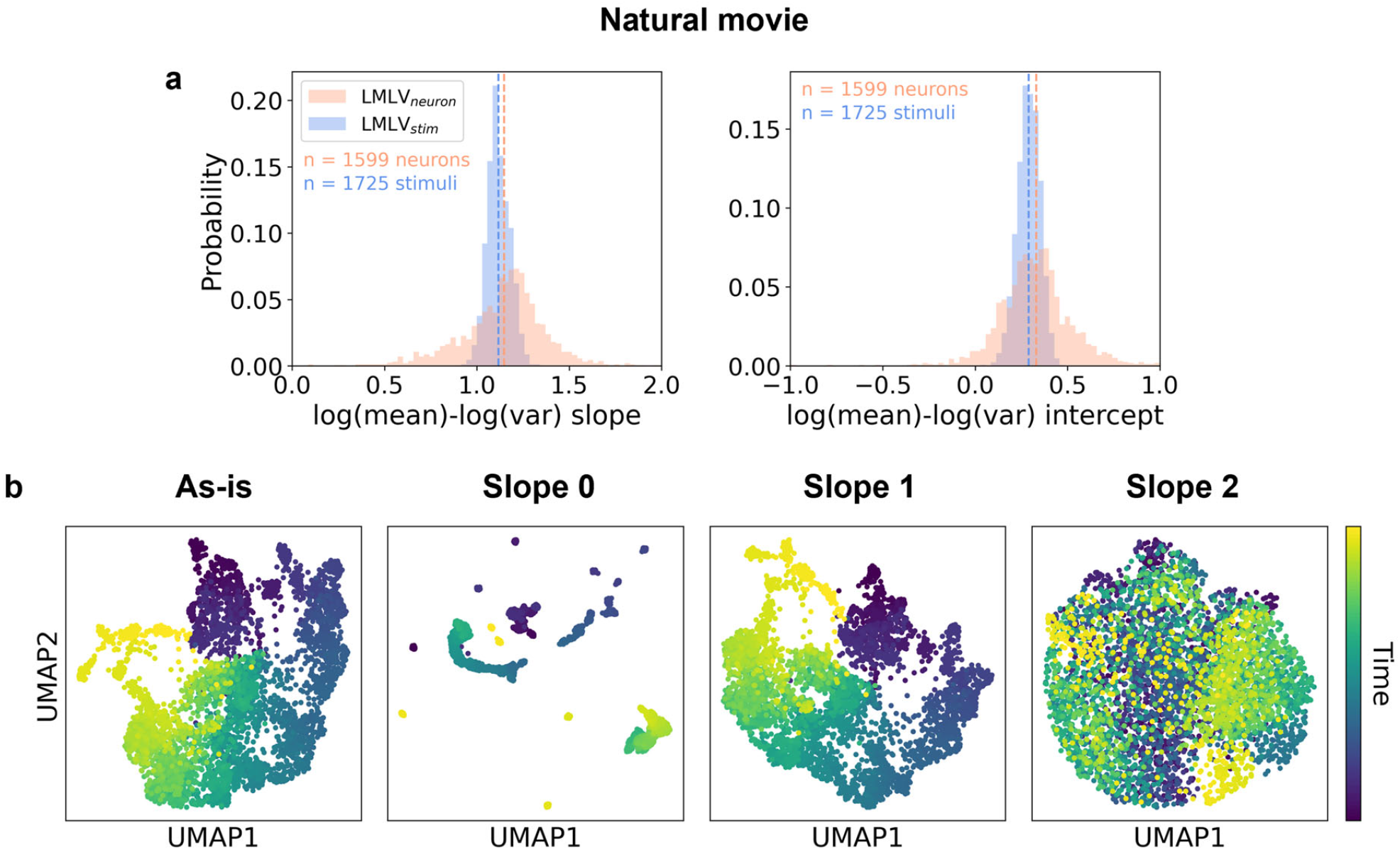
LMLV_stim_ slope 1 leads to continuous neural trajectories during a natural movie. **a** Histograms of LMLV_stim_ and LMLV_neuron_ slopes (*left*) and y-intercepts (*right*) across mouse V1 neurons’ responses to a natural movie (‘natural movie one’). Spike counts were calculated for 400 ms bins of a 30 s movie clip, resulting in 75 consecutive windows (‘stimuli’) repeated 60 times. Slopes and y-intercepts in all sessions were pooled (23 sessions of 23 mice). Dotted lines indicate the mean of each condition. **b** UMAP embedding of spike count vectors for a natural movie in an example session. Each dot corresponds to a repeat of a stimulus and color indicates the temporal order of the stimuli.

**Supplementary Fig. 5:**
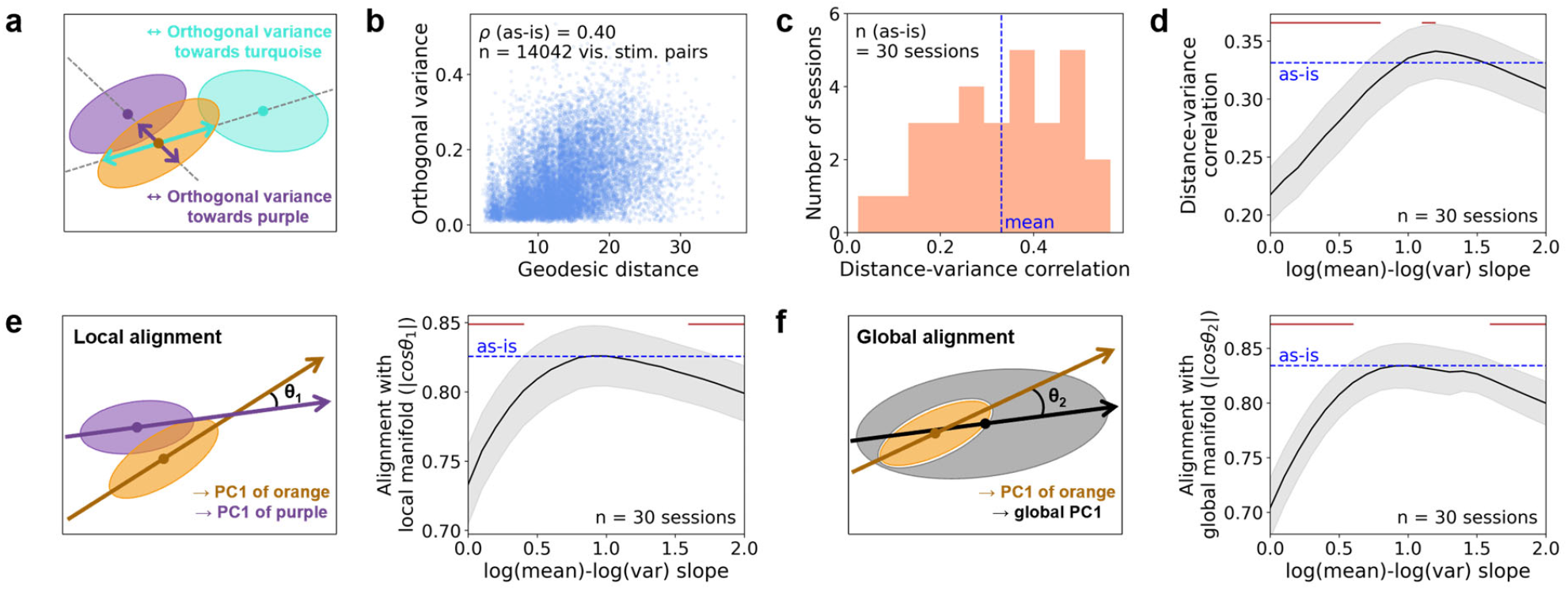
Geometry of trial-to-trial variability reflects the visual representation space. **a** Schematic illustration of orthogonal variance. Three stimulus manifolds are shown (*orange, purple* and *turquoise* ellipses) and the bold point inside each stimulus manifold is the centroid. **b** Scatterplot of the inter-centroid geodesic distance and orthogonal variance for the as-is data of an example session. The inter-centroid geodesic distance and the orthogonal variance between every pair of stimulus manifolds were plotted (119×118 = 14,042 pairs). Spearman correlation was denoted as ρ. **c** Histogram of Spearman correlations between inter-centroid geodesic distances and orthogonal variances. Spearman correlations were calculated as in **b** for each session. **d** Geodesic distance vs. orthogonal variance correlation. **e** Local alignment between neighboring pairs of stimulus manifolds. *Left*: Schematic illustration of two stimulus manifolds (*orange* and *purple* ellipses), where the *purple* manifold is the closest neighbor of the *orange* manifold. The local alignment of the *orange* manifold is defined as the absolute cosine similarity between PC1’s of the *orange* and *purple* manifolds (|cosθ_1_|). *Right*: Local alignment of 119 stimulus manifolds was averaged within each session. Local alignment peaked around LMLV_stim_ slope 1, indicating that neighboring stimulus manifolds were most aligned with each other at slope 1. **f** Global alignment of each stimulus manifold. *Left*: Schematic illustration of a stimulus manifold (*orange* ellipse) and all other stimulus manifolds (*gray*). The global alignment of the orange manifold is defined as the absolute cosine similarity between PC1’s of the *orange* and the remaining manifolds (|cosθ_2_|). *Right*: Global alignment of 119 stimulus manifolds within each session was averaged. Global alignment peaked around LMLV_stim_ slope 1, indicating that the shapes of individual stimulus manifolds most closely resembled the broader natural scenes representation space at slope 1. **d-f** Mean ± SEM across sessions was plotted as a function of LMLV_stim_ slopes. The red lines on top indicate slopes that are significantly different from the as-is data (p < 0.05, two-sided Wilcoxon signed-rank test across sessions, Holm-Bonferroni corrected).

**Supplementary Fig. 6:**
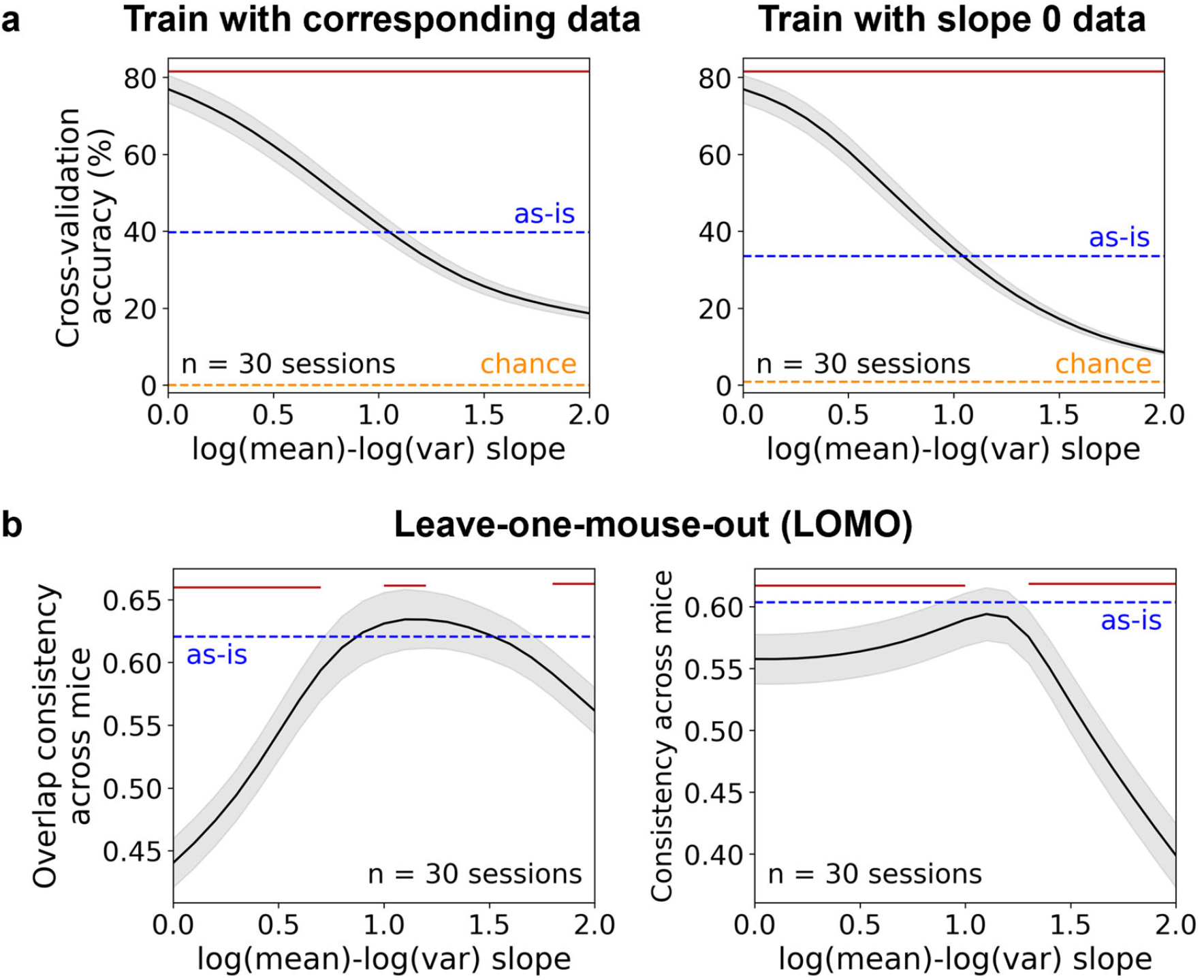
Cross-slope decoding accuracy and leave-one-mouse-out representational consistency. **a** 10-fold cross-validation classification accuracy of linear SVM decoder, trained within each LMLV_stim_ slope (*left*) and slope 0 (*right*). *Orange* dotted line indicates chance level (1/119 ≈ 0.008). **b** Overlap consistency (*left*) and consistency of RSMs (*right*) in leave-one-mouse-out (LOMO) analyses. Percentage of overlap was calculated for every pair of stimulus manifolds in each session. Both consistency metrics were calculated as the Spearman correlation between each session and the average of the other sessions. Representational consistency peaks around LMLV_stim_ slope 1, in agreement with pairs of mice (**Fig. 4d, e**). **a, b** Mean ± SEM across sessions was plotted as a function of LMLV_stim_ slopes. The red lines on top indicate slopes that are significantly different from the as-is data (p < 0.05, two-sided Wilcoxon signed-rank test across sessions, Holm-Bonferroni corrected).

**Supplementary Fig. 7:**
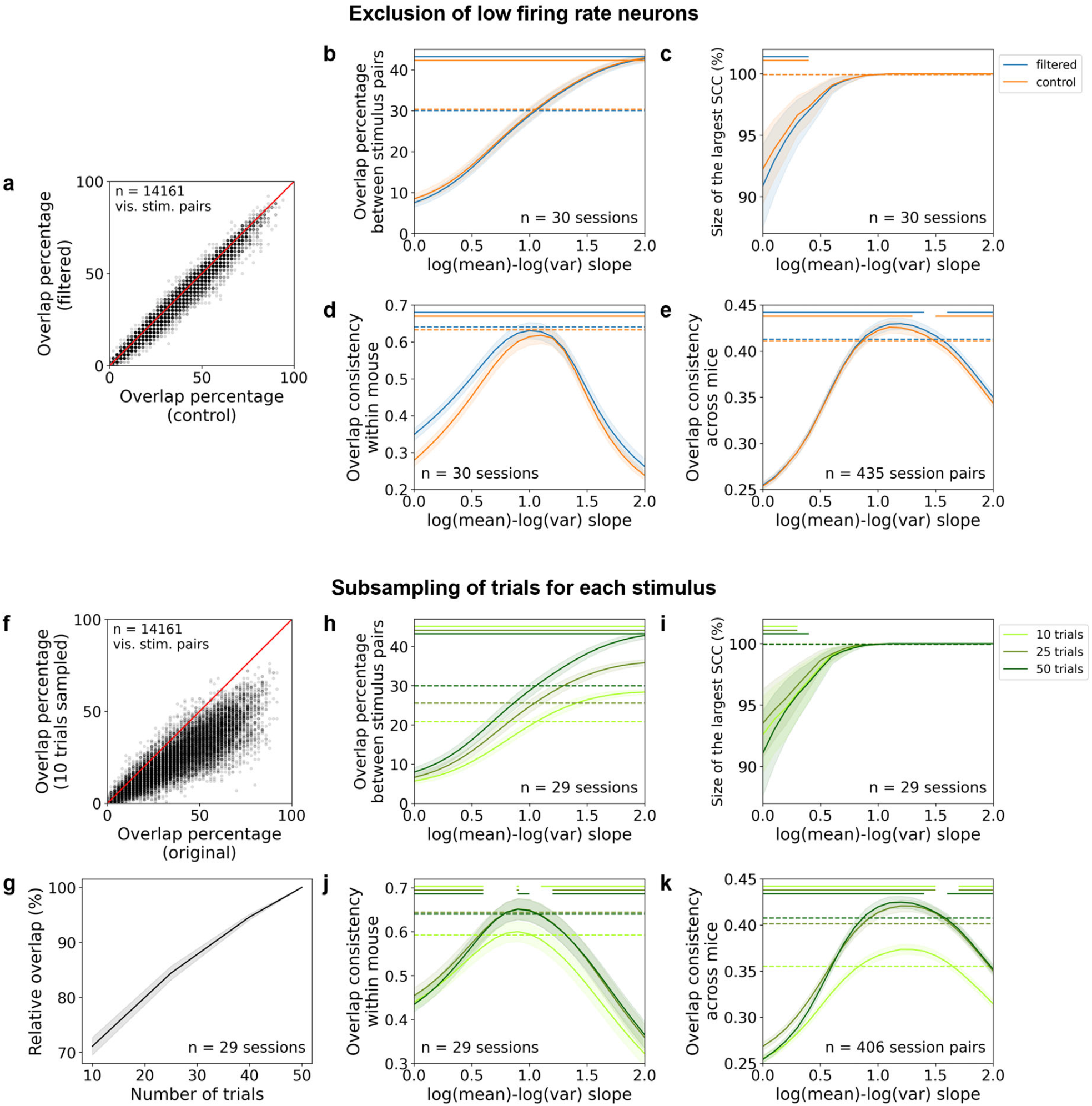
Relative overlap percentage is stable under neuron and trial subsampling. **a** Scatterplot of overlap percentage in an example session. In the filtered condition, we excluded low-firing rate neurons with mean spike count less than 1 across all stimuli. In the control condition, we randomly excluded the same number of neurons. **b** Overlap percentage in filtered and control conditions. **c** Size of the largest SCC in filtered and control conditions. **d** Overlap consistency between distinct neuronal subsets within each mouse in filtered and control conditions. **e** Overlap consistency between pairs of mice in filtered and control conditions. **f** Scatterplot of overlap percentage in an example session, between stimulus manifolds after randomly subsampling 10 trials from each stimulus, compared to all 50 trials per stimulus. **g** Relative overlap percentage between stimulus manifolds after subsampling trials from each stimulus. Overlap percentages were normalized by the condition with all repeats (50 trials). **h** Overlap percentage after subsampling trials from each stimulus. **i** Size of the largest SCC after subsampling trials from each stimulus. **j** Overlap consistency between distinct neuronal subsets within each mouse after subsampling trials from each stimulus. **k** Overlap consistency between pairs of mice after subsampling trials from each stimulus. **a, f** Overlap percentage was computed between every pair of stimulus manifolds in example sessions (119×119 = 14,161 pairs). **b-e, g-k** Mean ± SEM across sessions (**b-d, g-j**) or session pairs (**e, k**) was plotted as a function of LMLV_stim_ slopes. The colored lines on top of **b-e** and **h-k** indicate slopes that are significantly different from the as-is data (p < 0.05, two-sided Wilcoxon signed-rank test across sessions or session pairs, Holm-Bonferroni corrected). The dotted lines in **b-e** and **h-k** indicate the mean of as-is data for each condition. A session that has less than 50 trials per stimulus (47–50 trials) was excluded in **g-k**.

**Supplementary Fig. 8:**
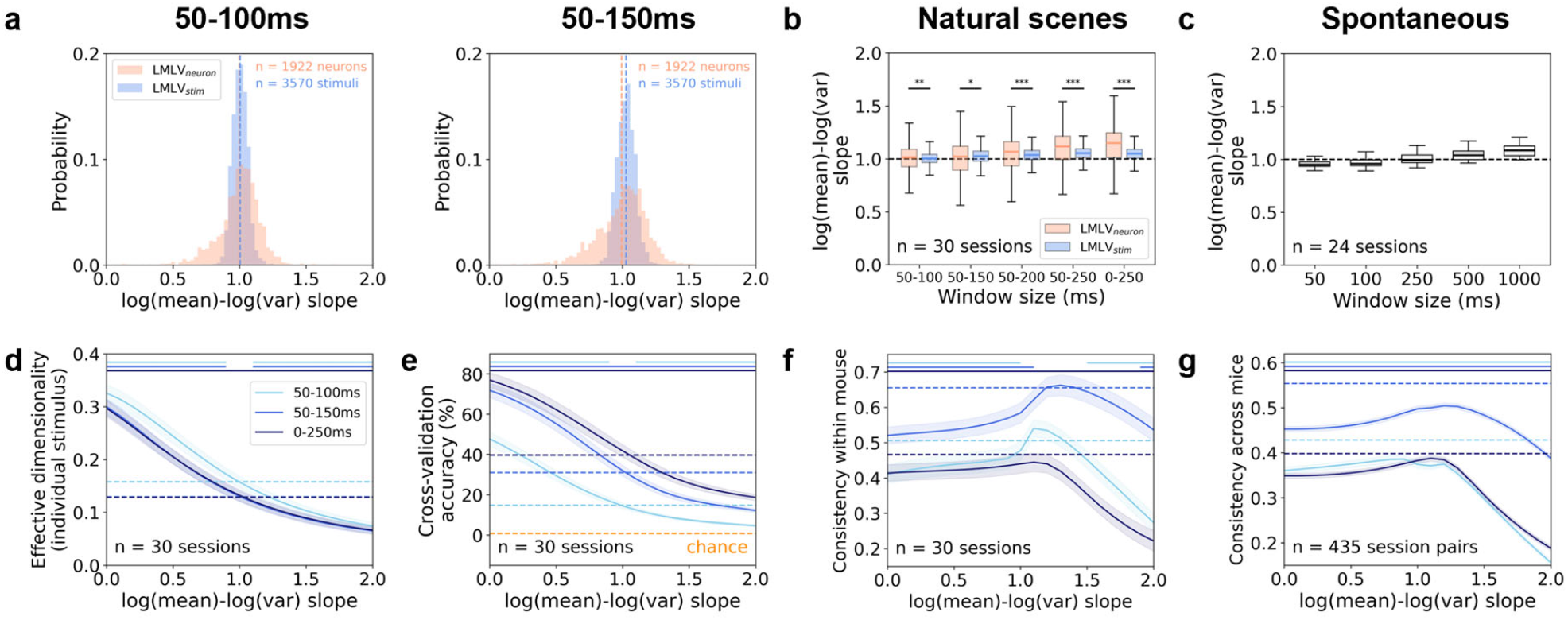
Results are consistent across spike-count time windows. **a** Histograms of LMLV_stim_ and LMLV_neuron_ slopes for spike counts of V1 neurons during 50–100 ms (*left*) and 50–150 ms (*right*) after natural scenes stimulus onset. Slopes in 30 sessions of 30 mice were pooled. The dotted lines indicate the mean of each condition. **b** Box plots of LMLV_stim_ and LMLV_neuron_ slopes using spike counts during different time windows. V1 neurons (LMLV_neuron_; n = 1,922) and natural scenes stimuli (LMLV_stim_; n = 3,570) were pooled across sessions, and median and IQR were plotted. Both LMLV_stim_ and LMLV_neuron_ slopes were significantly different across time windows (p < 10^-300^ for both LMLV_stim_ and LMLV_neuron_, Friedman test). The asterisks (*) on top indicate significant differences between LMLV_stim_ and LMLV_neuron_ (*p < 0.05, **p < 0.01, ***p < 0.001, two-sided Wilcoxon rank-sum test, Holm-Bonferroni corrected). **c** Box plots of log-mean vs. log-variance slopes across V1 neurons using spike counts during different time windows of a long spontaneous block. Long spontaneous blocks of 30 minutes were binned into different lengths of non-overlapping time windows. Median and IQR across sessions were plotted. The slopes were significantly different across time windows (p = 2.58×10^-19^, Friedman test). **d** Effective dimensionality of individual stimulus manifolds using spike counts during different time windows. **e** 10-fold cross-validation classification accuracy of linear SVM decoder using spike counts during different time windows. **f** Consistency of RSMs between distinct neuronal subsets within each mouse using spike counts during different time windows. **g** Consistency of RSMs between pairs of mice using spike counts during different time windows. **b, c** The dotted lines indicate the log-mean vs. log-variance slope of 1. **d-g** Mean ± SEM across sessions (**d-f**) or session pairs (**g**) was plotted as a function of LMLV_stim_ slopes. The colored lines on top indicate slopes that are significantly different from the as-is data (p < 0.05, two-sided Wilcoxon signed-rank test across sessions or session pairs, Holm-Bonferroni corrected). The dotted lines indicate the mean of as-is data for each time window.

**Supplementary Fig. 9:**
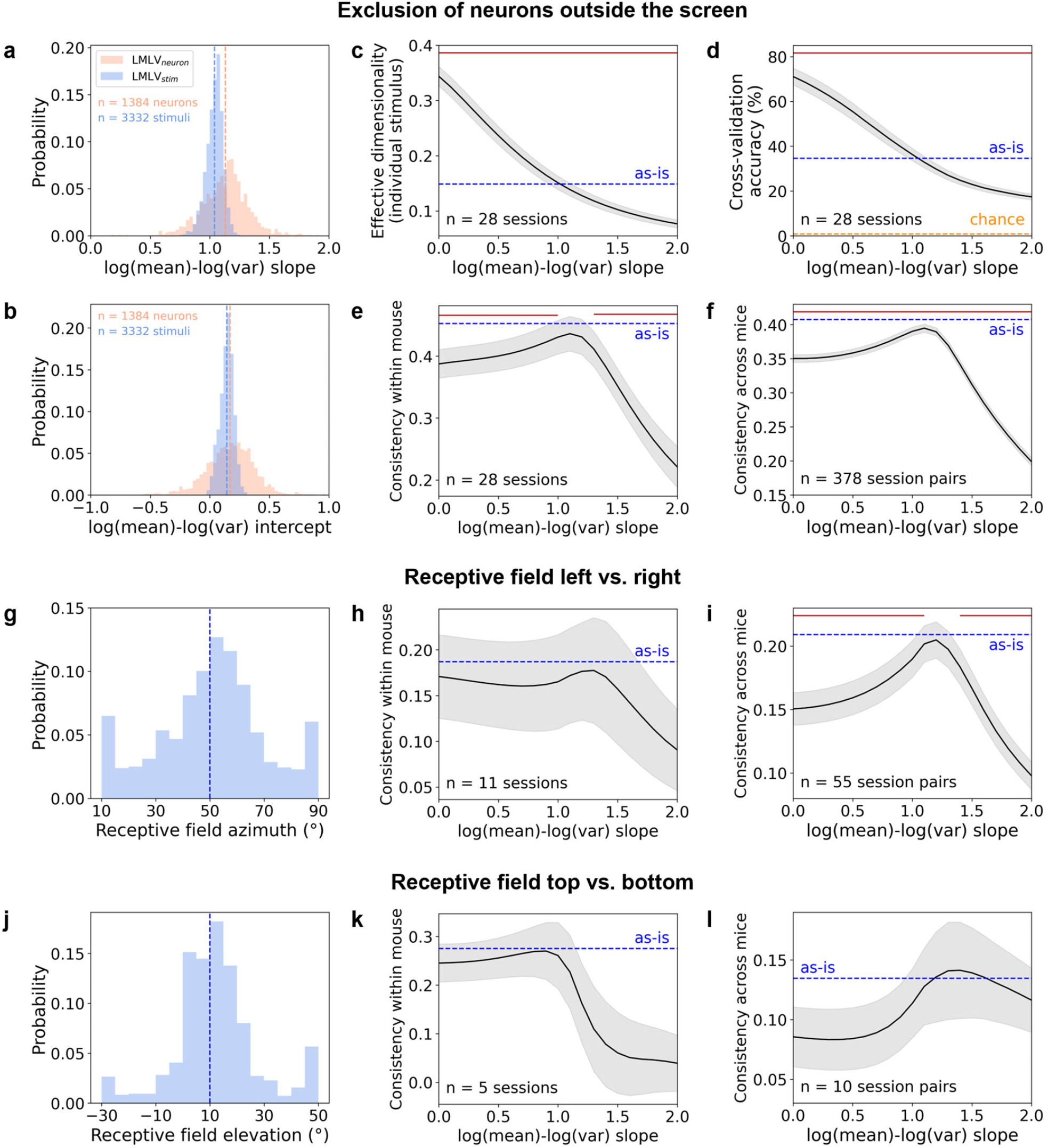
Results are consistent when accounting for receptive fields. **a, b** Histograms of LMLV_stim_ and LMLV_neuron_ slopes (**a**) and y-intercepts (**b**) on natural scene trials, using V1 neurons whose receptive fields were on the screen. The dotted lines indicate the mean of each condition. Sessions with fewer than 10 neurons on the screen were excluded, leaving 28 sessions from 28 mice. **c** Effective dimensionality of individual stimulus manifolds mice using V1 neurons whose receptive fields were on the screen. **d** 10-fold cross-validation classification accuracy of linear SVM decoder mice using V1 neurons whose receptive fields were on the screen. **e** Consistency of RSMs between distinct neuronal subsets within each mouse using V1 neurons whose receptive fields were on the screen. **f** Consistency of RSMs between pairs of mice using V1 neurons whose receptive fields were on the screen. **g** Histogram of V1 neuronal receptive field azimuth, pooling 30 sessions of 30 mice. The dotted lines indicate the center of azimuth used for neuron partitioning. **h, i** Consistency of RSMs between neuronal subsets with left vs. right receptive fields within each mouse (**h**) and between pairs of mice (**i**). **j** Histogram of V1 neuronal receptive field elevation, pooling 30 sessions of 30 mice. The dotted lines indicate the middle of elevation (**j**) used for neuron partitioning. **k, l** Consistency of RSMs between neuronal subsets with top vs. bottom receptive fields within each mouse (**k**) and between pairs of mice (**l**). **c-f, h, i, k, l** Mean ± SEM across sessions (**c-e, h, k**) or session pairs (**f, i, l**) was plotted as a function of LMLV_stim_ slopes on natural scenes trials. The *red* lines on top indicate slopes that are significantly different from the as-is data (p < 0.05, two-sided Wilcoxon signed-rank test across sessions or session pairs, Holm-Bonferroni corrected). Sessions with fewer than 10 neurons on the screen (**c-f**) or on each half of the screen (**h, i, k, l**) were excluded.

**Supplementary Fig. 10:**
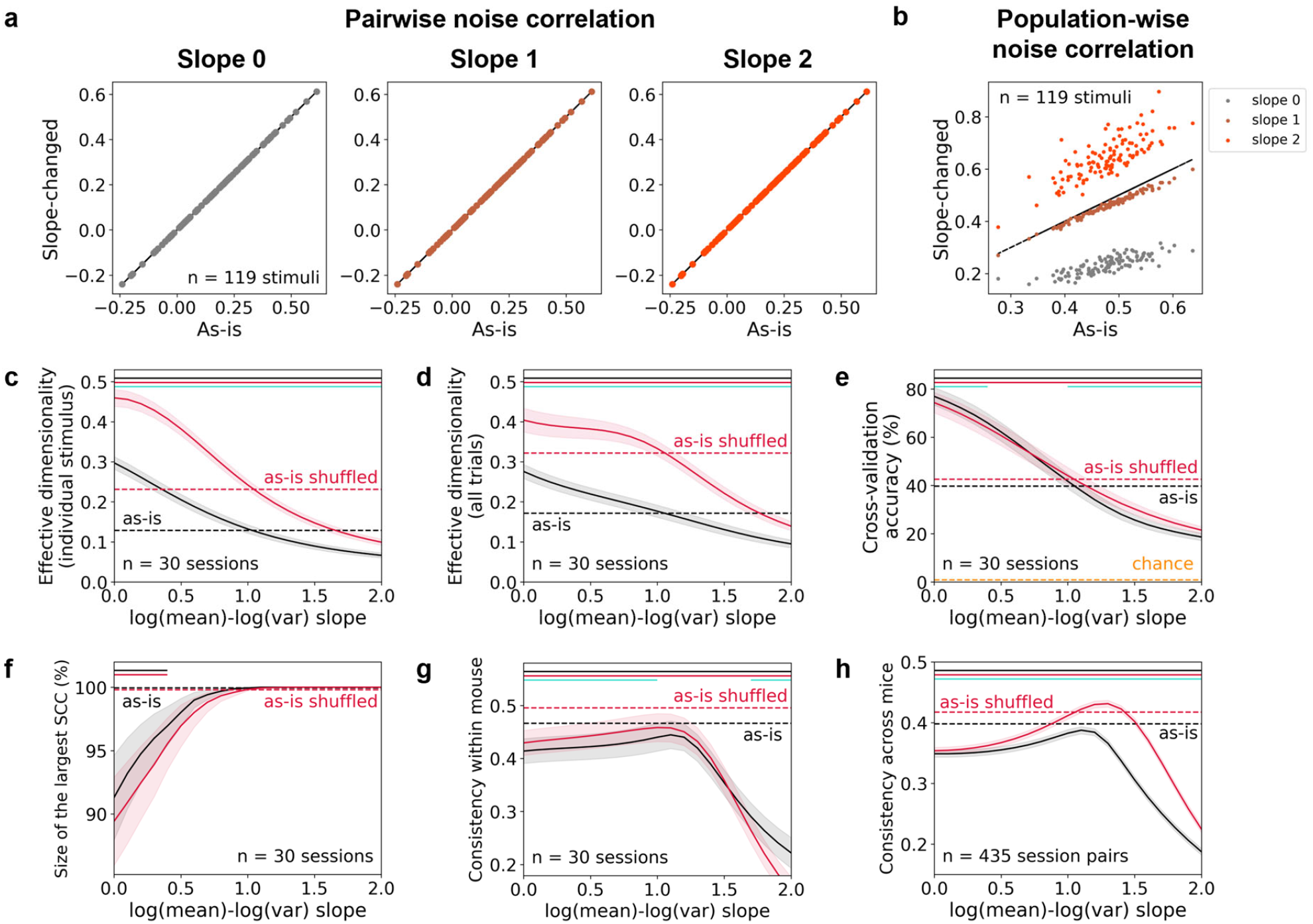
Interactions between LMLV_stim_ slope and noise correlations. **a** Stimulus-specific pairwise noise correlations of an example V1 neuron pair. **b** Stimulus-specific population-wise noise correlations, defined as the fraction of variance explained by PC1, for an example session. **a, b** Each dot corresponds to a stimulus and the *black* line indicates y=x. **c, d** Effective dimensionality of individual stimulus manifolds (**c**) and all stimulus manifolds (**d**), for shuffled (*red*) and original (*black*) trial order. Shuffling randomized the trial order within a stimulus for each neuron, such that pairwise noise correlations were destroyed. **e** 10-fold cross-validation classification accuracy of linear SVM decoder for shuffled (*red*) and original (*black*) trial order. **f** Size of the largest SCC for shuffled (*red*) and original (*black*) trial order. **g** Consistency of RSMs between distinct neuronal subsets within each mouse for shuffled (*red*) and original (*black*) trial order. **h** Consistency of RSMs between pairs of mice for shuffled (*red*) and original (*black*) trial order. **c-h** Mean ± SEM across sessions (**c-g**) or session pairs (**h**) was plotted as a function of LMLV_stim_ slopes. The *red* and *black* lines on top indicate slopes that are significantly different from spike counts without slope manipulation, in shuffled (‘as-is shuffled’) and original (‘as-is’) trial order, respectively. The *turquoise* lines on top indicate slopes where the shuffled and original trials are significantly different. All three lines indicate p < 0.05, two-sided Wilcoxon signed-rank test across sessions or session pairs, Holm-Bonferroni corrected.

**Supplementary Fig. 11:**
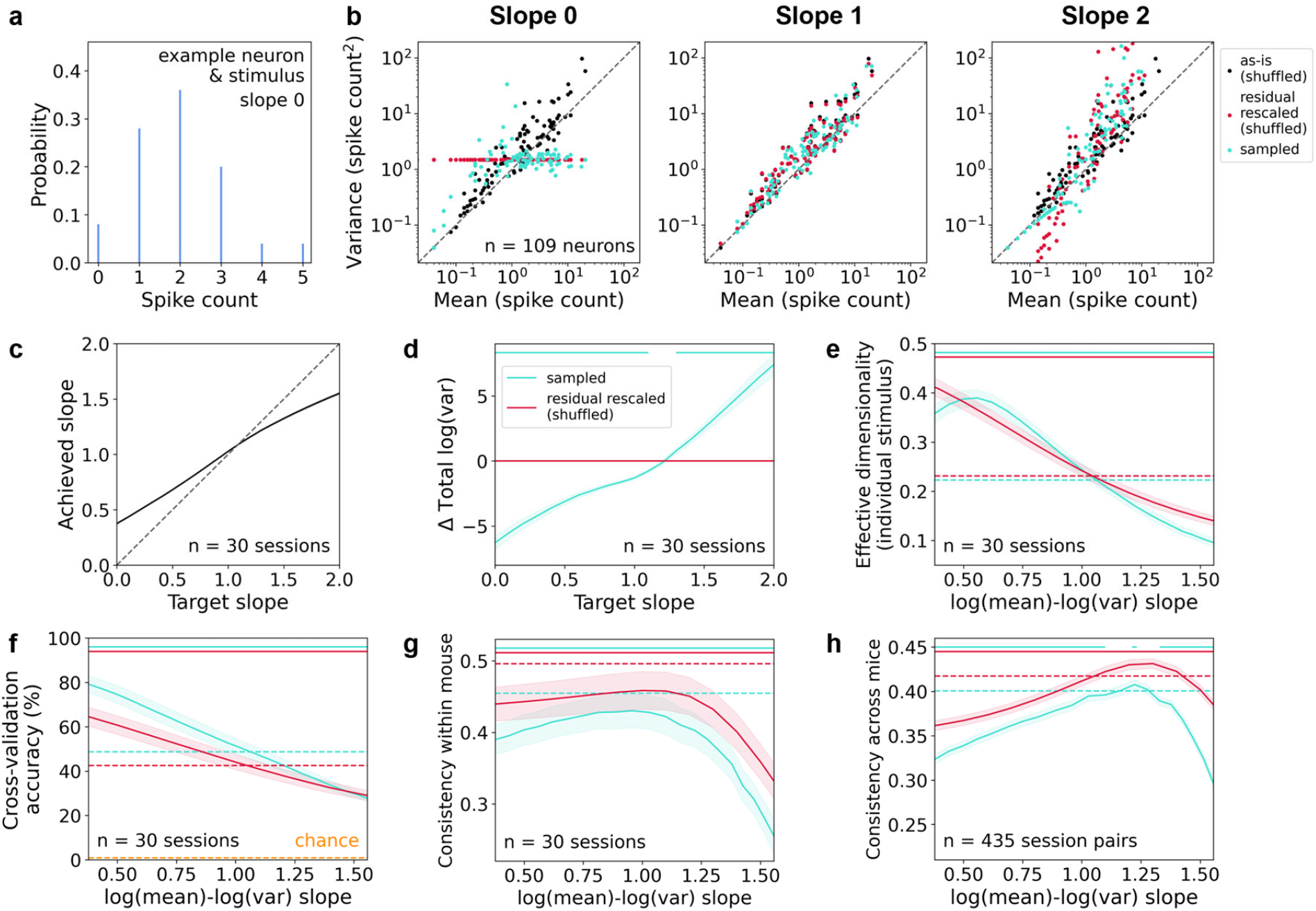
LMLV_stim_ slope manipulation via spike count sampling from discrete probability distributions is consistent with the residual rescaling method. **a** Spike count distribution sampled for an example neuron-stimulus combination. Spike counts were sampled from negative binomial (variance ≥ 1.05×mean), Poisson (0.95×mean < variance < 1.05×mean), or binomial (variance ≤ 0.95×mean), depending on the target dispersion. The sampling was independent across neurons, eliminating pairwise noise correlations. **b** LMLV_stim_ scatterplots of an example session and a natural scene stimulus. The *black* dots denote as-is data (no slope manipulation); *red* dots denote residual-rescaled pseudo-data; and the *turquoise* dots denote spike counts sampled from discrete probability distributions (negative binomial, Poisson, or binomial distribution). **c** LMLV_stim_ slopes of sampled spike counts. Slopes were averaged across stimuli within each session. **d** Total log-variance of sampled spike counts and residual-rescaled data relative to as-is data. Log-variance was summed across neurons within each stimulus (total log-variance), and the difference from as-is (slope-manipulated data – as-is data) was averaged across stimuli within each session. **e** Effective dimensionality of individual stimulus manifolds for sampled spike counts (*turquoise*) and residual-rescaled data (*red*). **f** 10-fold cross-validation classification accuracy of linear SVM decoder for sampled spike counts (*turquoise*) and residual-rescaled data (*red*). **g** Consistency of RSMs between distinct neuronal subsets within each mouse for sampled spike counts (*turquoise*) and residual-rescaled data (*red*). **h** Consistency of RSMs between pairs of mice for sampled spike counts (*turquoise*) and residual-rescaled data (*red*). **b-h** Trial order in as-is and residual-rescaled data were shuffled to eliminate noise correlations, to compare with the sampling method where spike counts for each neuron and stimulus were sampled independently. Mean ± SEM across sessions (**c-g**) or session pairs (**h**) was plotted as a function of LMLV_stim_ slopes. The *turquoise* and *red* lines on top of **d-h** indicate slopes that are significantly different between slope-manipulated and original-slope data, for the spike count sampling (*turquoise*) and residual rescaling (*red*) methods, respectively. Both lines indicate p < 0.05, two-sided Wilcoxon signed-rank test across sessions or session pairs, Holm-Bonferroni corrected. In **e-h**, the x-axis denotes achieved LMLV_stim_ slopes (**c**).

**Supplementary Fig. 12:**
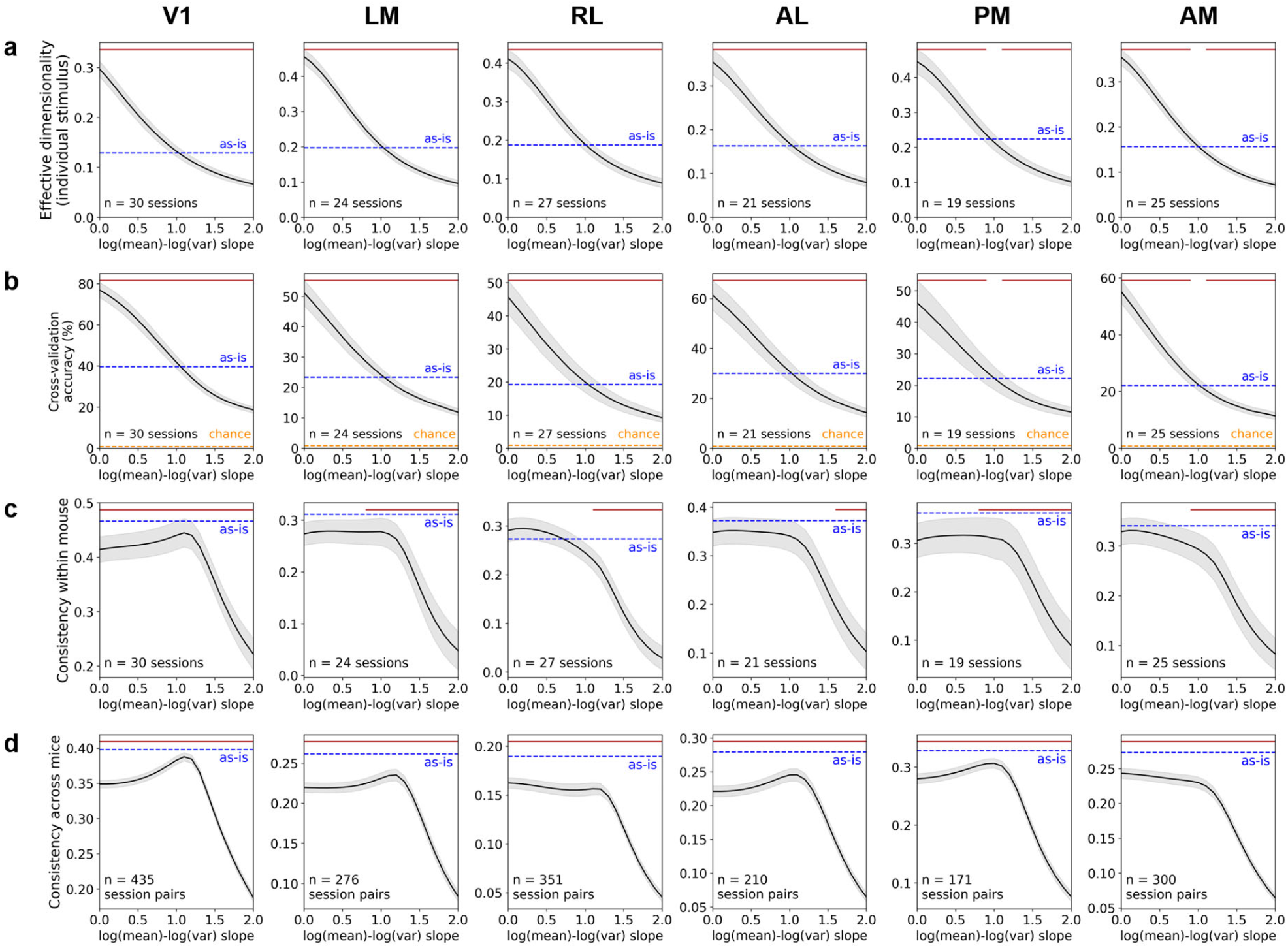
Dimensionality, decoding and representational similarity analyses for higher visual cortices. **a** Effective dimensionality of individual stimulus manifolds. **b** 10-fold cross-validation classification accuracy of the linear SVM decoder. **c** Consistency of RSMs between distinct neuronal subsets within each mouse. **d** Consistency of RSMs between pairs of mice. **a-d** Mean ± SEM across sessions (**a-c**) or session pairs (**d**) was plotted as a function of LMLV_stim_ slopes for 6 visual cortical areas. The *red* lines on top indicate slopes that are significantly different from the as-is data (p < 0.05, two-sided Wilcoxon signed-rank test across sessions or session pairs, Holm-Bonferroni corrected).

**Supplementary Fig. 13:**
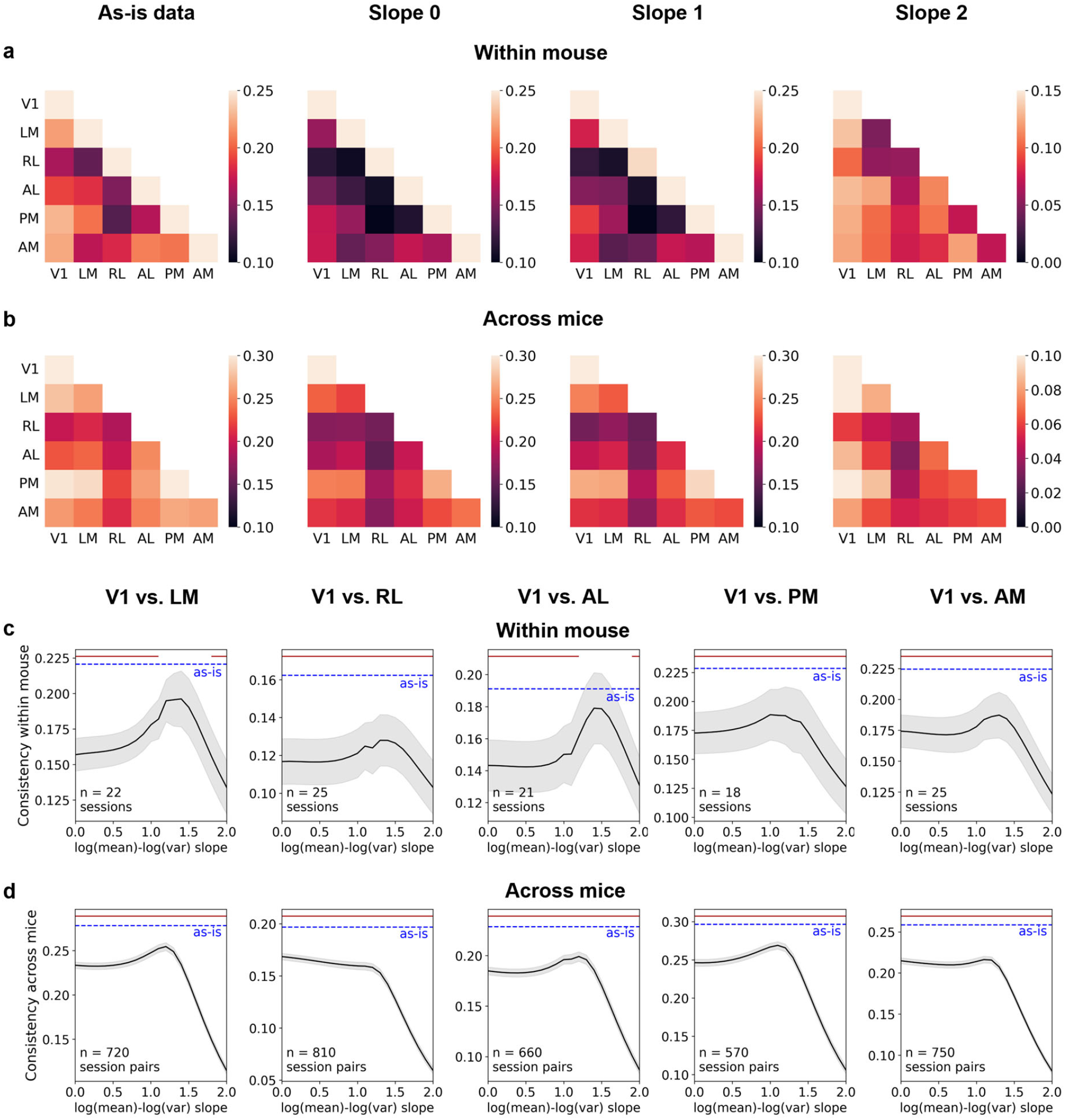
Inter-area representational similarity analyses. **a, b** Heatmaps of the consistency of RSMs between distinct visual cortical areas within each mouse (**a**) and between pairs of mice (**b**). Mean across sessions (**a**) or session pairs (**b**) was plotted. **c, d** Consistency of RSMs between V1 and higher visual areas within each mouse (**c**) and between pairs of mice (**d**). Mean ± SEM across sessions (**c**) or session pairs (**d**) was plotted as a function of LMLV_stim_ slopes. The *red* lines on top indicate slopes that are significantly different from the as-is data (p < 0.05, two-sided Wilcoxon signed-rank test across sessions or session pairs, Holm-Bonferroni corrected). **a-d** LMLV_stim_ slopes were fit and manipulated within each visual cortical area. For within-mouse analyses (**a, c**), we randomly sampled neurons from each of the two areas, using a sample size equal to half the number of neurons in the area with fewer neurons.

**Supplementary Fig. 14:**
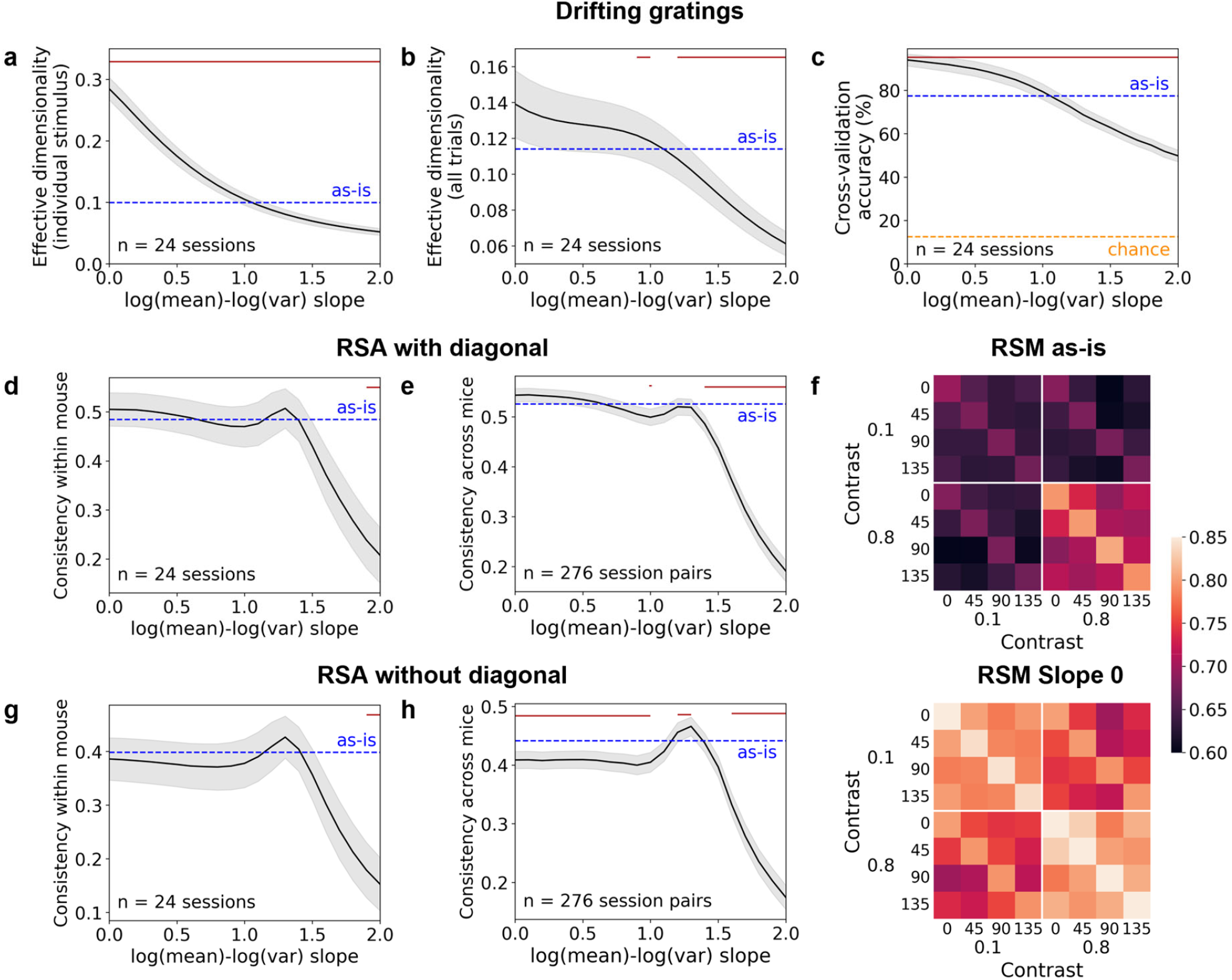
Dimensionality, decoding and representational similarity analyses for drifting gratings. **a, b** Effective dimensionality of individual stimulus manifolds (**a**) and all stimulus manifolds (**b**) for drifting gratings. Spike counts during 0–250 ms after stimulus onset were used. 8 drifting gratings were presented, in 4 directions, 2 contrasts, and a fixed temporal frequency of 2 Hz. Each drifting grating was repeated 75 times in 24 sessions of 24 mice. **c** 10-fold cross-validation classification accuracy for the linear SVM decoder of drifting gratings. **d** Consistency of RSMs between distinct neuronal subsets within each mouse for drifting gratings. **e** Consistency of RSMs between pairs of mice for drifting gratings. **f** Heatmaps of RSMs for drifting gratings averaged across sessions. **g** Consistency of RSMs excluding the diagonals, between distinct neuronal subsets within each mouse for drifting gratings. **h** Consistency of RSMs excluding the diagonals, between pairs of mice for drifting gratings. **a-e, g, h** Mean ± SEM across sessions (**a-d, g**) or session pairs (**e, h**) was plotted as a function of LMLV_stim_ slopes. The *red* lines on top indicate slopes that are significantly different from the as-is data (p < 0.05, two-sided Wilcoxon signed-rank test across sessions or session pairs, Holm-Bonferroni corrected).

**Supplementary Fig. 15:**
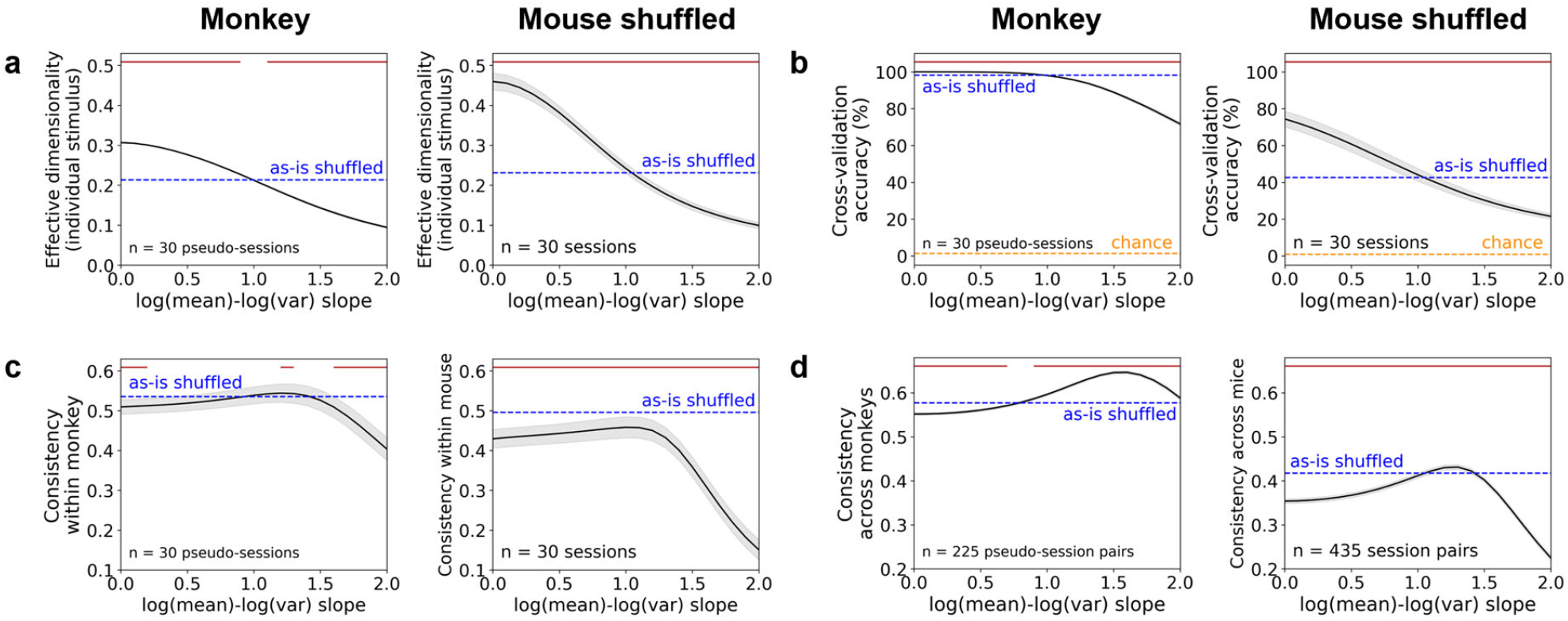
Slope manipulation effects on natural scenes representation are largely consistent between mouse and monkey V1. **a** Effective dimensionality of individual stimulus manifolds in monkeys (*left*) and mice (*right*). **b** 10-fold cross-validation classification accuracy of the linear SVM decoder in monkeys (*left*) and mice (*right*). **c** Consistency of RSMs between distinct neuronal subsets within each subject in monkeys (*left*) and mice (*right*). **d** Consistency of RSMs between different subjects in monkeys (*left*) and mice (*right*). **a-d** For monkey data, V1 single unit spike counts during 40–160 ms after stimulus onset were analyzed, where 75 natural scenes were repeated ∼40 times in 32 sessions of 2 monkeys (238 V1 neurons in 15 sessions of monkey M1, 220 V1 neurons in 17 sessions of monkey M2). In each monkey, neurons were pooled across sessions and subsampled into 15 pseudo-sessions of 110 neurons (total 30 pseudo-sessions). In addition, we bootstrapped 50 trials for each stimulus, and trial order within each stimulus was shuffled for each neuron to remove noise correlations. The same shuffling procedure was applied to mouse V1 natural scenes data. Mean ± SEM across sessions (**a-c**) or session pairs (**d**) were plotted as a function of LMLV_stim_ slopes. *Red* lines on top indicate slopes that are significantly different from the as-is shuffled data in *blue* (p < 0.05, two-sided Wilcoxon signed-rank test across sessions or session pairs, Holm-Bonferroni corrected).

**Supplementary Fig. 16:**
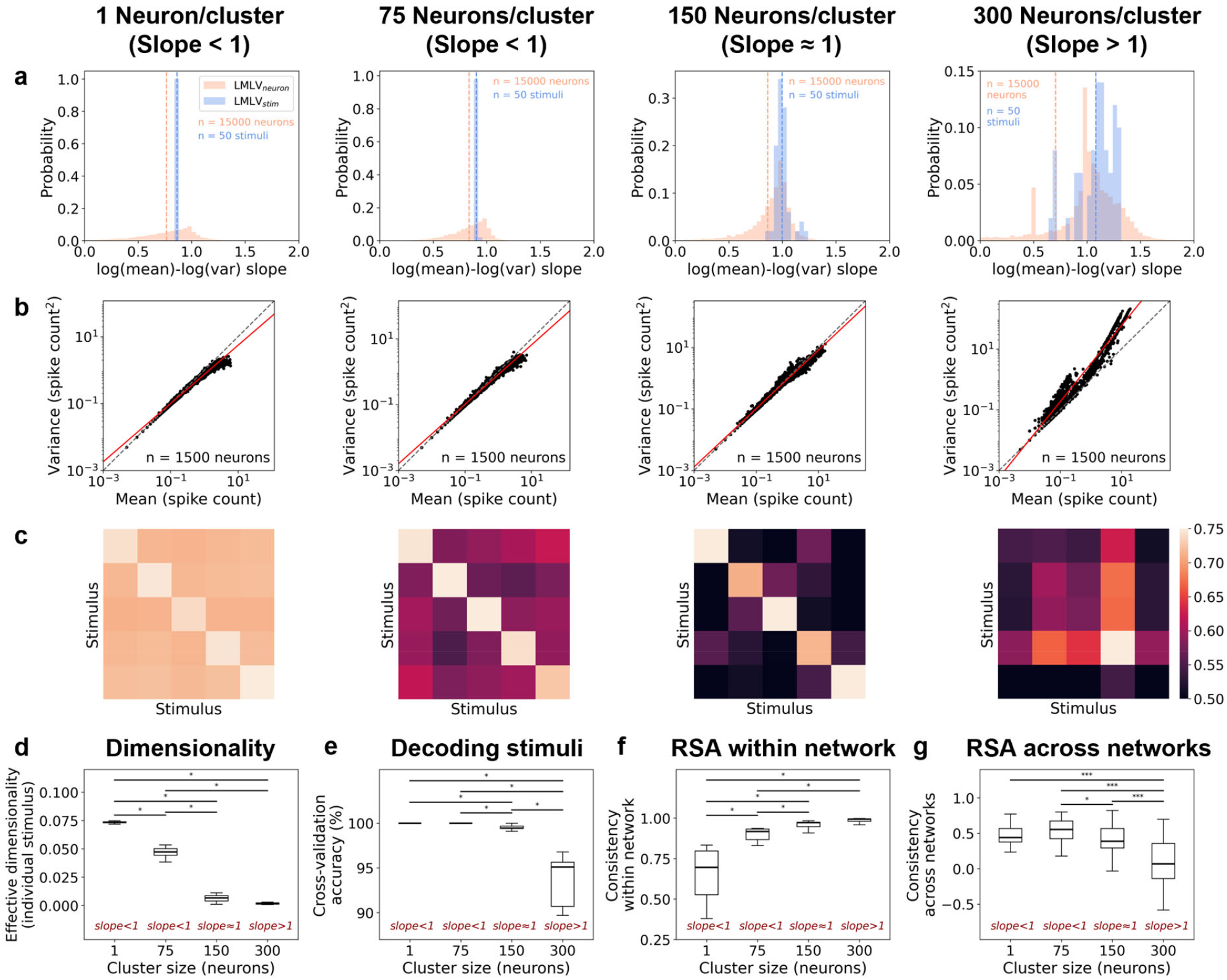
Dimensionality, decoding and representational similarity analyses in neural network simulations. **a** Histograms of LMLV_stim_ and LMLV_neuron_ slopes for clustered recurrent network simulations. Slopes in 10 networks with the same cluster size were pooled. The dotted lines indicate the mean of each condition. **b** LMLV_stim_ scatterplots of an example network and a stimulus. The *red* lines are the lines fit by linear regression of the *black* dots. **c** RSMs of example networks with different cluster sizes. **d** Effective dimensionality of individual stimulus manifolds for clustered recurrent networks. **e** 10-fold cross-validation classification accuracy of the linear SVM decoder for clustered recurrent networks. **f** Consistency of RSMs between distinct neuronal subsets within each network. **g** Consistency of RSMs between pairs of networks. **d-g** Median and IQR across 10 networks (**d-f**) or 45 network pairs (**g**) were plotted. The asterisks (*) on top indicate significant differences (*p < 0.05, **p < 0.01, ***p < 0.001, two-sided Wilcoxon signed-rank test across networks or network pairs, Holm-Bonferroni corrected).

